# Emergence of curvature-dependent tissue growth from cell crowding and mechanical interactions

**DOI:** 10.64898/2026.03.10.710423

**Authors:** Shahak Kuba, Matthew J. Simpson, Pascal R. Buenzli

## Abstract

Many biological tissues grow at rates that depend strongly on the local geometry of the tissue interface, yet the cellular mechanisms underlying this dependence remain poorly understood. In this work, we develop a two-dimensional mathematical model of tissue growth to investigate how curvature-dependent growth emerges from the complex interactions between cell-scale crowding and mechanical interactions. The tissue interface is represented as a confluent layer of mechanically interacting cells that simultaneously produce new tissue. We formally derive a continuum limit of this discrete model, obtaining a system of partial differential equations governing the coupled evolution of cell density and tissue geometry. Although curvature is not explicitly represented in the discrete model, curvature-dependent tissue growth emerges naturally in the continuum limit through geometric crowding associated with tissue production. The continuum equations also establish a direct relationship between cellular mechanical properties and the effective diffusivity governing mechanical relaxation at the tissue scale. Numerical simulations illustrate that the competition between tissue production and mechanical relaxation can result in a variety of tissue evolutions, including tissue interface smoothing consistent with curvature-controlled *in vitro* growth in porous scaffolds and bone formation. Finally, the model predicts that the time required for scaffold pore closure scales with pore size but also depends on a pore shape factor, extending previous empirical observations of linear scaling with size. More broadly, this work provides an analytical framework linking cell-scale mechanics to tissue-scale growth and geometry.

## 1 Introduction

Biological tissue growth during development and regeneration is governed by complex interactions between tissue geometry and tissue mechanics (Rodriguez et al., 1994; Taber, 1995; Jones and Chapman, 2012; Goriely, 2017; Ambrosi et al., 2019). Cellular growth induces local cell crowding and mechanical forces that influence the evolution of tissue geometry. Mechanical stresses may also directly regulate cellular activities such as cell proliferation, migration, and differentiation through mechanobiological processes, affecting both growth and geometry (Chen et al., 2004; Nelson et al., 2005; Discher et al., 2005; Wang and Thampatty, 2006; Lim et al., 2010; Eyckmans et al., 2011; Jansen et al., 2015; Ladoux and Mège, 2017; Xi et al., 2019). The complexity of these spatiotemporal interactions make it challenging to understand how tissue shape develops during growth, and how tissue geometry influences growth through its effects on cellular crowding and mechanical stress. Understanding these coupled processes is of particular importance in tissue engineering and regenerative medicine, where controlling tissue growth is essential for regenerating damaged or diseased tissues (Hollister, 2005; O’Brien, 2011; Bidan et al., 2013; Dzobo et al., 2018).

Several experimental studies have shown that the local curvature of the tissue interface can strongly influence the rate at which the tissue grows (Rumpler et al., 2008; Dunlop et al., 2010; Bidan et al., 2012, 2013, 2016; Fratzl et al., 2023; Buenzli et al., 2020; Lanaro et al., 2021; Schamberger et al., 2023) (Figure 1). Figure 1a shows an *in-vitro* experiment of tissue growth within a 3D-printed square-shaped scaffold pore. Initially, cells grow near the pore corners, where curvature is highest. Growth subsequently remains faster in these regions than along the pore edges, resulting in a substantially thicker tissue layer in the corners after 14 days. These observations suggest that the curvature of the local tissue interface influences the speed of the tissue interface in a similar way to mean curvature flow, where interface velocity is directly proportional to curvature (Grayson, 1987; Crandall and Lions, 1983). Similar observations are made *in-vivo*, such as in bone tissues, where new bone formation tends to smoothen the bone surface through curvature-dependent growth (Wozniak and El Haj, 2007; Ripamonti and Roden, 2010; Alias and Buenzli, 2018) (Figure 1b).

**Figure 1.**
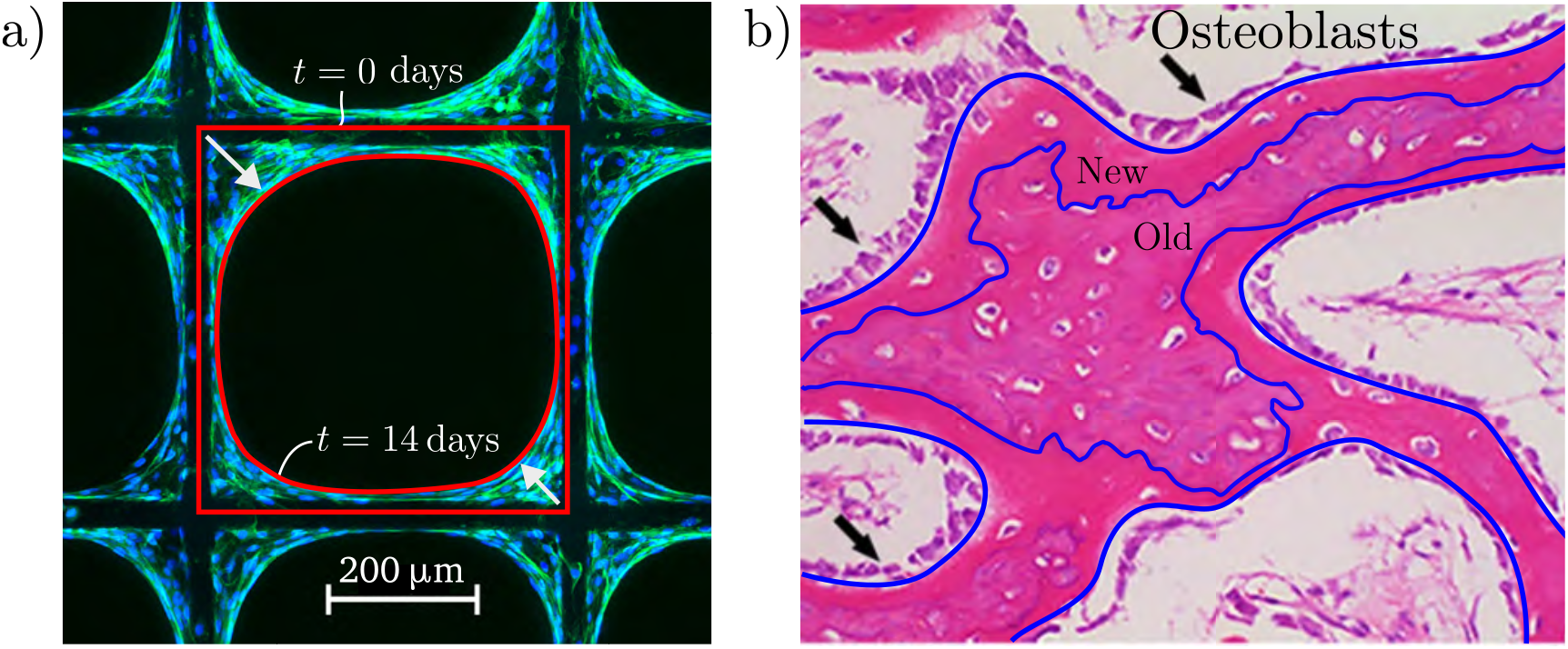
Experimental images of *in vitro* and *in vivo* tissue growth. (a) Cell culture experiment within a porous scaffold, courtesy of Brenna Devlin and Maria Woodruff. The image, taken 14 days into the experiment, highlights the tissue interface in red at both *t* = 0 and *t* = 14 days, demonstrating the smoothing behaviour associated with curvature-controlled tissue growth. (b) Hematoxylin and eosin staining shows osteoblasts along the edge of bone trabeculae in contact with each other. The tissue interface, highlighted in blue, illustrates bone matrix development at early and late stages of bone formation behind a series of osteoblasts. Reproduced with permissions from Han et al. (2016).

Curvature-dependent growth has been hypothesised to arise in part due to crowding effects and mechanics-induced relaxation (Alias and Buenzli, 2017, 2018, 2019; Hegarty-Cremer et al., 2021; Buenzli et al., 2020; Buenzli and Simpson, 2022). However, no mathematical framework currently exists that couples curvature-induced cell crowding, mechanical cell relaxation, and tissue growth within a unified model. The aim of this paper is to develop such a framework and determine how the complex spatio-temporal coupling between cellular-scale crowding effects and mechanical relaxation gives rise to curvature-dependent growth at the tissue scale.

Previous mathematical models of biological tissue growth include morphoelastic theories based on continuum mechanics (Rodriguez et al., 1994; Dunlop et al., 2010; Jones and Chapman, 2012; Goriely, 2017; Ambrosi et al., 2019; Taber, 2020; Schamberger et al., 2023). These models describe growth through phenomenological growth tensors that are difficult to connect to individual cell properties. Similarly, mean curvature flow models, in which the normal velocity of the tissue interface is proportional to its curvature, are not explicitly related to cellular quantities (Rumpler et al., 2008; Bidan et al., 2012; Guyot et al., 2014). Existing cell-based mechanical models are either one-dimensional (Murray et al., 2009; Fozard et al., 2010; Baker et al., 2019; Murphy et al., 2019; Tambyah et al., 2020), or rely primarily on numerical simulations (Bi et al., 2015; Osborne et al., 2017; Ghaffarizadeh et al., 2018; Kim et al., 2021; Vetter et al., 2024), limiting our understanding of how curvature-induced growth emerges from cellular interactions using first principles.

In this paper, we develop a mathematical model for tissue growth in which the tissue interface is represented by a confluent layer of individual cells interacting mechanically like mechanical springs. New tissue production displaces these cells in the normal direction, resulting in a coupled evolution of cell density and interface geometry. The discrete cell formulation explicitly tracks the trajectories and stress state of individual cells. By accounting for the motion of the interface in two-dimensional space through new tissue production, this model builds significantly on the model of mechanical interactions on a static substrate proposed in (Buenzli et al., 2025). A nontrivial feature of our model is the prescription of microscopic evolution rules by which neighbouring cells with different orientations and progressing in different spatial directions, must interact mechanically to obtain biologically realistic tissue evolutions. We formally derive a continuum limit of this discrete cell model, leading to a system of partial differential equations for the coupled evolution of cell density and tissue interface.

A significant outcome of our analysis is that cellular-scale crowding and mechanical relaxation lead to the emergence of curvature-controlled tissue growth and nonlinear cell diffusion in the continuum limit. The continuum evolution equations for cell density and tissue interface obtained in the limit can be directly related to those of the (non-mechanical) continuum models of Alias and Buenzli (2017, 2018, 2019) and Hegarty-Cremer et al. (2021). Curvature explicitly arises from crowding, and cell diffusivity is expressed in terms of the mechanical properties of individual cells. Numerical simulations of the model show that the evolution of the tissue depends significantly on the competition between growth and mechanical relaxation. Finally, our mathematical model predicts that scaffold pore closure time in tissue engineering constructs scales with pore size but depends on a geometric shape factor, thereby generalising the linear scaling reported in Buenzli et al. (2020).

## 2 Model description

Our mathematical model describes biological tissues that primarily grow due to new tissue production near the tissue surface. This is a common growth pattern observed in many biological tissues such as bone, where osteoblasts on the bone surface secrete new bone matrix (Figure 1b) (Wozniak and El Haj, 2007; Ripamonti and Roden, 2010), tumour spheroids, and tissues grown in porous scaffolds, where cells proliferate and generate new extracellular matrix within a narrow band close to tissue surface (Figure 1a) (Rumpler et al., 2008; Bidan et al., 2012, 2013, 2016; Guyot et al., 2014; Lee et al., 2019; Buenzli et al., 2020; Lanaro et al., 2021; Riccobelli, 2025).

We first present a discrete population model of cells that interact mechanically and generate new tissue material (Section 2.1), before formally deriving a continuum limit of this model, leading to a continuum model governed by PDEs (Section 2.2). A list of symbols used in the discrete and continuum models is provided in Table 1.

**Table 1.** Mathematical symbols of the discrete and continuum models. See Section 2.4 for further detail on parameter values or ranges.

| Symbol | Description | Value(s)/Units |
| --- | --- | --- |
| **Discrete model** |  |  |
| $k$ | Mechanical spring stiffness | $10\text{--}10^3 \text{ N } \mu\text{m}^{-1}$ |
| $a$ | Resting spring length | $1\text{--}3 \text{ } \mu\text{m}$ |
| $\eta$ | Drag coefficient | $1 \text{ N day } \mu\text{m}^{-1}$ |
| $k_f$ | Tissue formation rate per cell | $87.74 \text{ } \mu\text{m}^2 \text{ day}^{-1}$ |
| $N$ | Number of cells | - |
| $m$ | Number of springs per cell | $1\text{--}10$ |
| $\ell_i$ | Spring length | $\mu\text{m}$ |
| $\rho_i$ | Spring density | $\mu\text{m}^{-1}$ |
| $q_i$ | Cell density | $\mu\text{m}^{-1}$ |
| **Continuum model** |  |  |
| $k_f$ | Tissue formation rate per cell | $87.74 \text{ } \mu\text{m}^2 \text{ day}^{-1}$ |
| $D$ | Diffusivity | $1\text{--}10^5 \text{ } \mu\text{m}^2 \text{ day}^{-1}$ |
| $\ell$ | Spring length | $\mu\text{m}$ |
| $s$ | Arc length | $\mu\text{m}$ |
| $\rho$ | Spring density | $\mu\text{m}^{-1}$ |
| $q$ | Cell density | $\mu\text{m}^{-1}$ |
| $\kappa$ | Curvature | $\mu\text{m}^{-1}$ |

### 2.1 Discrete model

Similarly to Buenzli et al. (2025), we represent the curved tissue interface as a chain of *N* connected cells (Figure 2a), where each cell body is represented by *m* sub-cellular components. These components are assumed to interact mechanically like springs, allowing us to represent both cell body deformations and cell mechanics. For simplicity, we will refer to these sub-cellular components as *springs* in the remainder of the paper. The *i*^th^ spring of the tissue interface is represented by the two-dimensional position vectors of its boundaries ***r***_*i*−1_(*t*) and ***r***_*i*_ (*t*) at time *t*, for *i* = 1, 2, …, *M*, where *M* = *mN* is the total number of springs on the tissue interface. We choose the interface to be closed, like in the pore closing experiment of Figure 1a, such that ***r***_0_(*t*) = ***r***_*M*_ (*t*) at all times.

**Figure 2.**
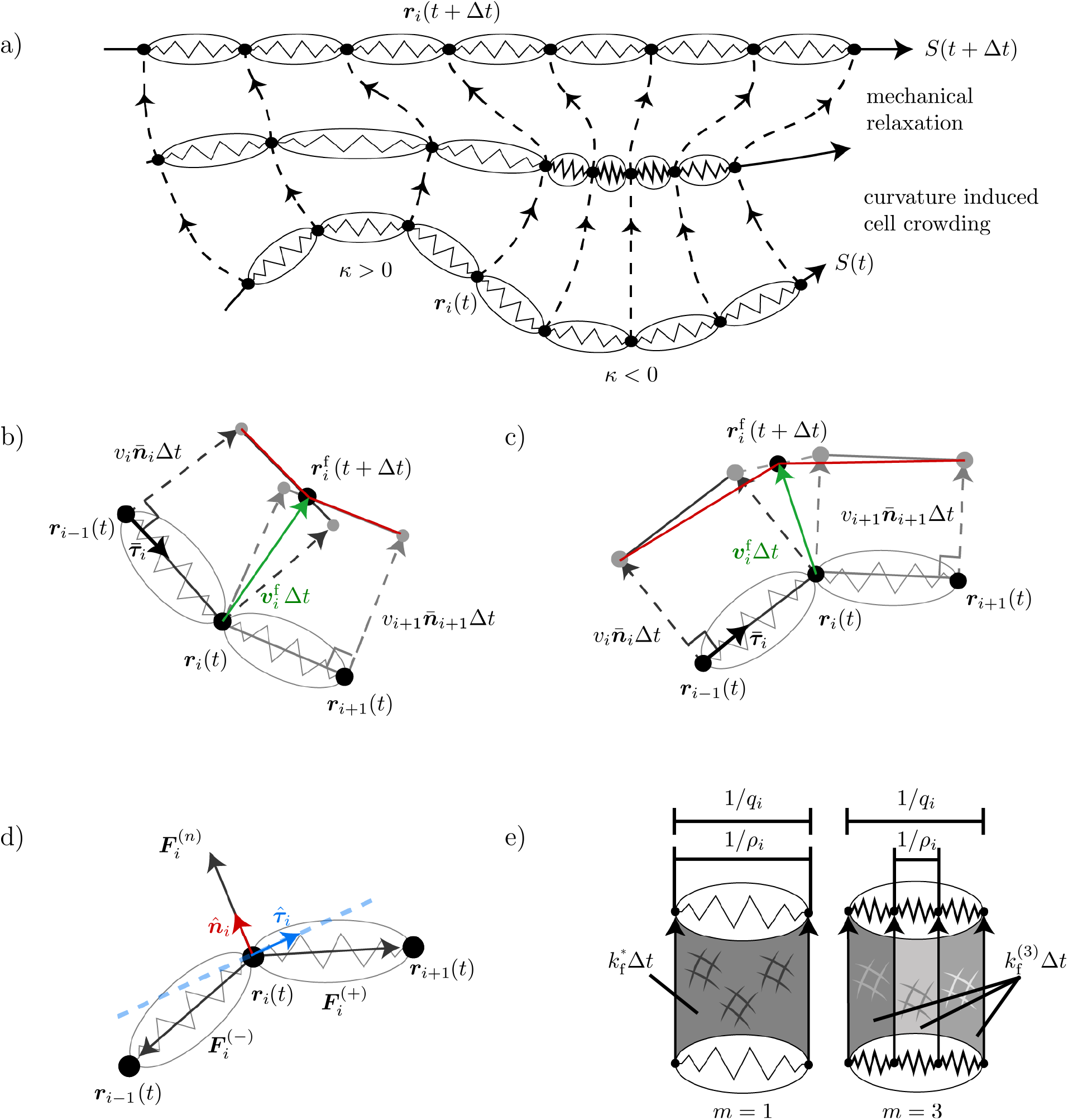
Discrete model of cells. (a) Evolution of the tissue interface over a time step *Δt*. The interface is represented by a polygonal boundary *S t* defined by a chain of cells each containing *m* springs (*m* = 1 in the figure for simplicity). During a time increment *Δt*, cells deposit new tissue and are displaced perpendicularly to the interface *S* (*t*). Cells crowd where the interface is concave (*κ* < 0), and spread out where the interface is convex (*κ* > 0). Crowding is countered by mechanical relaxation, which tends to redistribute the cells along the new interface *S* (*t + Δt*). Although depicted sequentially, these two processes occur simultaneously during *Δt*. (b)-(c) The velocity 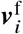 (green) of spring boundary *i* is determined by its new boundary position 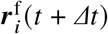 When neighbouring springs evolve toward each other and overlap, 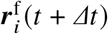 is the intersection of the normally displaced segments of each spring, resulting in compressed springs (red, (b)). When neighbouring springs move away from each other, 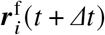 is set as the average position of the middle spring boundaries after their normal displacement, resulting in elongated springs (red, (c)). (d) Spring restoring forces 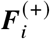 and 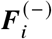, and normal reaction force 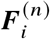 of the tissue substrate exerting at ***r***_*i*_, resulting in a net force along 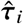. (e) The area of new tissue produced per spring per unit time 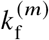 must be scaled with *m* so that two cells of equal length 1/*q*_*i*_ but comprising different numbers of springs deposit the same area of tissue per unit time 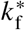.

We assume that cells at the tissue interface generate new tissue material at rate *k*_f_ (tissue area formed per cell per unit time). This new tissue material may comprise both extracellular matrix and embedded cells. In the continuum cell population models of Buenzli (2015); Alias and Buenzli (2017); Hegarty-Cremer et al. (2021), this mode of tissue growth evolves the tissue interface with a normal velocity given by

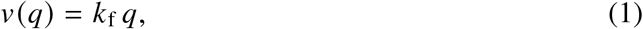

where *q* is the cell density (cells per unit length of the interface). In these continuum models, cell density is in turn affected by the evolution of the interface due to crowding effects. In our discrete model, however, the evolution of the tissue interface and of the cell density is determined by how spring boundaries evolve. We decompose the evolution of spring boundaries based on a velocity contribution 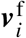 due to new tissue formation, and a velocity contribution 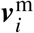 due to mechanical interaction between the springs:

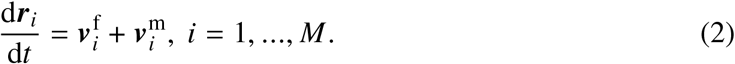

The following sections describe how we model the velocities 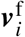 and 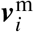.

#### 2.1.1 Formation of new tissue material

Because spring boundaries are vertices of a polygonal interface, it is not straightforward to specify how these vertices should evolve to model the normal velocity in Equation (1) prescribed in continuum frameworks. We define the growth-induced velocity 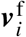 at time *t* in the discrete model by assuming a fixed time increment Δ*t* and set

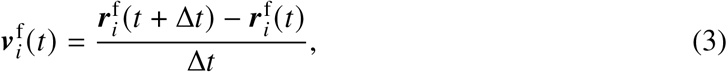

where 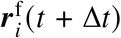 is the new position of the spring boundary after a time step Δ*t*. This new position is defined algorithmically as follows:

1. First, each spring *i* is displaced individually in its normal direction 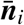 by a distance *v*_*i*_Δ*t*, where *v*_*i*_ is a normal velocity that depends on cell density by analogy with Equation (1) (Figure 2b,c). The normal vector 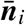 is the outward-pointing vector perpendicular to

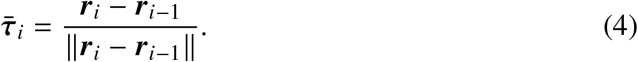
2. If the two springs adjacent to node *i* are on a concave portion of the interface, their normal displacement makes them intersect (Figure 2b). In this case, 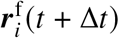 is defined as the intersection point, giving

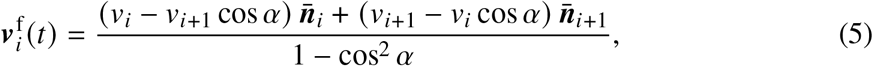

where *α* is the angle between the two adjacent springs, i.e.,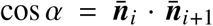. This microscopic evolution rule results in shorter springs, representing crowding.
3. If the two springs adjacent to node *i* are on a convex portion of the interface, their normal displacement makes them move away from each other, and they are no longer connected in their new configuration (Figure 2c). In this case, we define 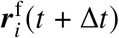 as the average position of the two evolved inner spring boundaries, giving

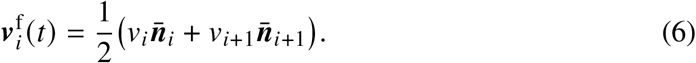

This evolution rule results in more elongated springs, representing spreading or dilution. The definition of 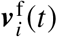 in Equations (5)–(6) is carried out without considering changes in spring lengths, since mechanical relaxation is represented separately by 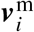. Numerically, this is similar to discretising Equation (2) using an operator splitting scheme: the contribution due to tissue formation can be calculated by assuming there is no mechanical relaxation, and the contribution due to mechanics in Section 2.1.2 can be calculated by assuming there is no interface motion. These assumptions are exact in the limit Δ*t* → 0.

#### 2.1.2 Mechanical relaxation

To define the velocity 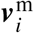 due to mechanical relaxation we consider the sum of forces exerting on the spring boundaries and follow the works of Murray et al. (2009); Fozard et al. (2010); Murphy et al. (2019); Buenzli et al. (2025). Each spring boundary ***r***_*i*_ experiences a force 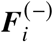 due to spring *i*, a force 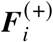 due to spring *i* + 1, and a normal reaction force 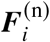 from the tissue substrate (Figure 2d) (Buenzli et al., 2025). We assume that cells evolve in a viscous medium, such that all inertial effects are negligible and their motion is overdamped. In the overdamped regime, the velocity contribution of the *i*th spring boundary due to mechanical relaxation is given by

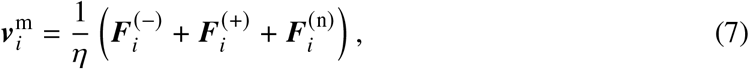

where *η* > 0 is the viscous drag coefficient. The spring restoring forces 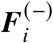 and 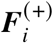 are given by 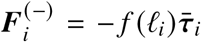 and 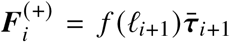, where *f* (*ℓ*) is a restoring force law defined as a function of the spring length *ℓ*. The normal reaction force 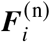 is chosen to be exactly opposite to the component of 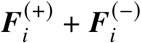 along the direction 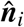 perpendicular to

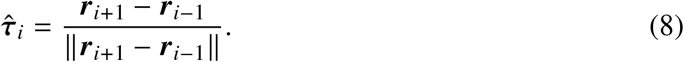

This choice of 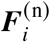 means that mechanical relaxation occurs only in the direction of the interface. With these assumptions, Equation (7) becomes:

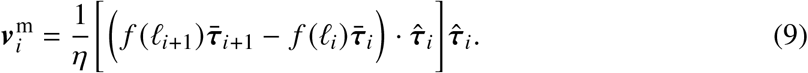

We will consider two distinct restoring force laws: the Hookean restoring force, given by

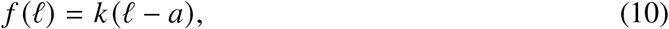

and a nonlinear restoring force law, given by

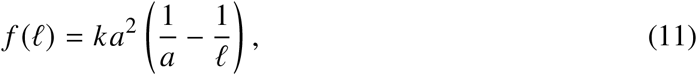

where *k* > 0 represents the spring stiffness, and *a* > 0 is the resting spring length. The nonlinear restoring force is such that *f* (*ℓ*) saturates for large elongations, limiting the interaction strength between widely separated cells, and diverges as *ℓ* → 0, preventing cells from collapsing to zero size, as assumed in Buenzli et al. (2025). Experimental measurement of interaction forces suggest similar behaviours (Gardel et al., 2004; Storm et al., 2005). For small displacements about the resting length *a*, both force laws are identical. As in Buenzli et al. (2025), the tangential stress *σ*_*ττ*_ in a subcellular component can be defined as

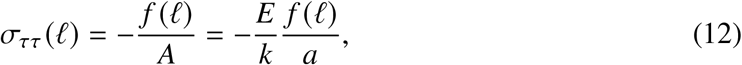

where *E* is the Young’s modulus of the cell body and *A* is the cell’s cross sectional area. Negative values of *σ*_*ττ*_ are associated with cells in extension.

### 2.2 Continuum limit

To describe the collective evolution of the discrete model of cells at a broader tissue scale, we derive a continuum limit of the discrete model defined by Equation (2). We follow the derivation of Buenzli et al. (2025) and expand cell densities about small spring lengths such that for a constant number of cells *N*, the number of springs per cell *m* becomes large, i.e., *m* → ∞. By expanding cell density about small spring lengths, we ensure that in the limit, cell density remains finite as spring density diverges.

The evolution of spring density *ρ*_*i*_ = 1/*ℓ*_*i*_ in the discrete model is given by

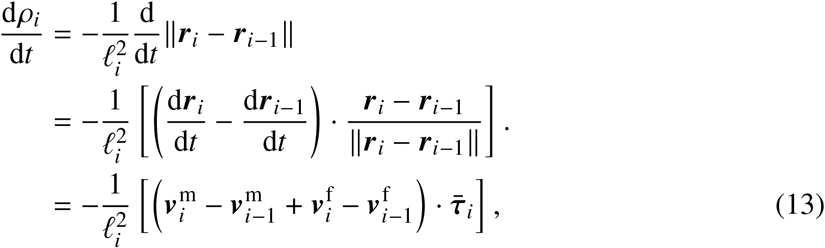

where in the last step we used Equations (2) and (4). As *m* becomes large, the tissue interface approaches a smooth curve. We represent this interface by a time-dependent curve ***r*** (*s, t*) parametrised by arc length *s* with 0 ≤ *s* ≤ *L* (*t*), where *L* (*t*) is the total length of the tissue interface at time *t*. The position ***r***_*i*_ (*t*) of a spring boundary can be referred to by its arc-length coordinate *s*_*i*_ (*t*), such that

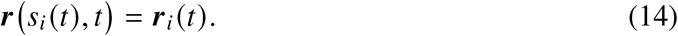

Similarly, we introduce a continuous spring density *ρ*(*s, t*) of arc length *s* and time *t*, such that

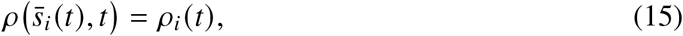

where 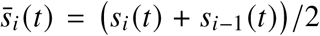 follows the arc-length coordinate of the midpoint of the *i*th spring (Buenzli et al., 2025). In the following, we omit explicit time dependences of arc-length positions to simplify notation, and write *s*_*i*_ and 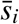 instead of *s*_*i*_ (*t*) and 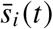.

To evaluate the right hand side of Equation (13) as *m* becomes large, we first note that the length of a spring becomes *ℓ*_*i*_ ~ *s*_*i*_ − *s*_*i*−1_ (Buenzli et al., 2025), and that the vectors 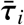 and 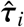 in Equations (4) and (8) both approach the unit tangent vector of ***r*** (*s, t*) at *s*_*i*_:

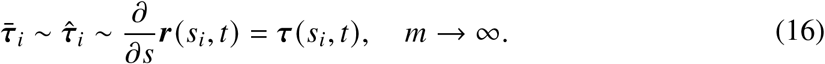

Similarly, 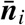 and 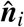 approach the outward unit normal vector ***n***(*s, t*) of ***r*** (*s, t*) at *s*_*i*_:

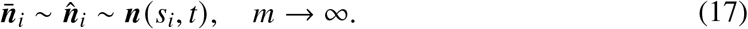

Furthermore, adjacent springs become parallel so that 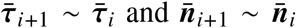. The velocity contribution 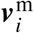 in Equation (9) thus becomes

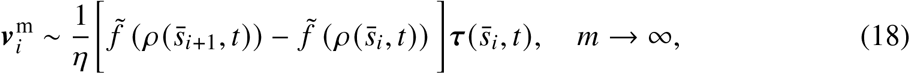

where 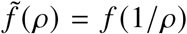 expresses the restoring force law as a function of spring density instead of spring length.

Because adjacent springs become parallel as *m* → ∞, Equation (5) for the velocity 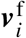 becomes identical to the average 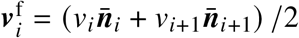 in Equation (6). This can be seen by taking *α* → 0 and *v*_*i*+1_ ~ *v*_*i*_ in Equation (5). Therefore, the change in local density due to new tissue production in Equation (13) becomes, as *m* → ∞,

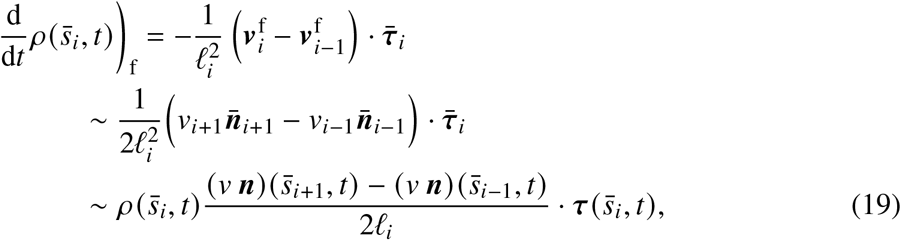

where 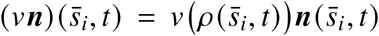, and *v* (*ρ*) is a continuous velocity function such that 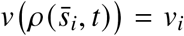 is the normal velocity of the *i*^th^ spring, and ***n***(*s, t*) is the unit normal vector to the interface in the continuum limit. For the last line in Equation (19), we used the fact that 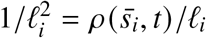. Substituting Equations (18) and (19) into Equation (13) gives, as *m* → ∞,

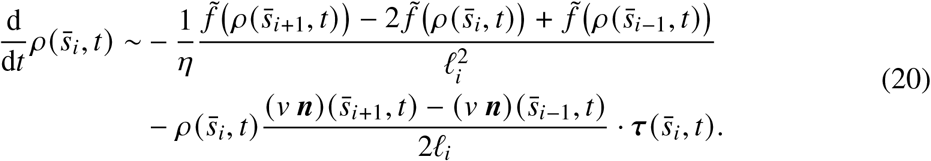

Differentiating the time dependence of 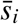 on the left hand side gives

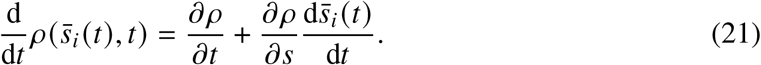

The evolution of arc-length positions 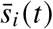 can be obtained by differentiating Equation (14) with respect to *t* and using Equation (2):

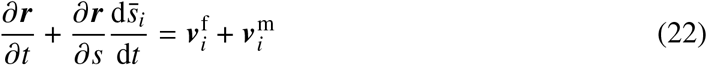

Since *∂* ***r***/*∂s* = ***τ*** is the unit tangent vector to the interface and 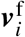 is perpendicular to ***τ***, projecting Equation (22) onto ***τ*** gives

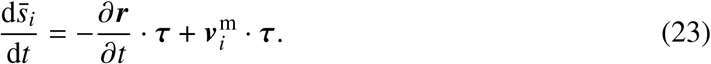

Substituting Equation (23) into Equation (21), the left hand side of Equation (20) becomes

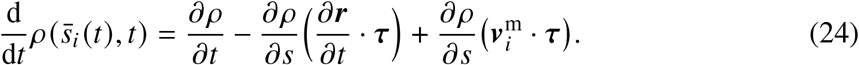

While the partial derivative *∂ ρ*/*∂t* represents changes in *ρ* following lines of constant arc length *s*, the first two terms on the right hand side of Equation (24) represent changes in *ρ* following normal trajectories to the interface, expressed in arc-length parametrisation ***r*** (*s, t*) (Alias and Buenzli, 2017; Hegarty-Cremer et al., 2021). We denote this contribution by

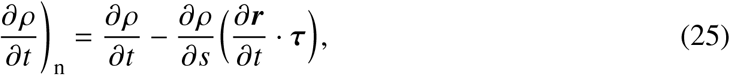

as in Hegarty-Cremer et al. (2021). As shown in Appendix A, the last term in the right hand side of Equation (24) cancels out precisely with some terms arising from expanding the first line on the right hand side of Equation (20) in a truncated Taylor series about 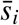 as *l*_*i*_ → 0. This cancellation is the same as that shown in Buenzli et al. (2025) on a static interface. Similar Taylor expansions carried out on the second line of Equation (20) show that (see Appendix A), as *m* → ∞,

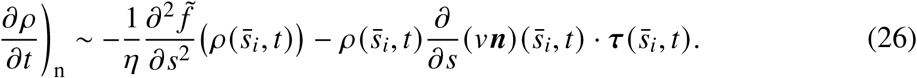

The last term on the right hand side of Equation (26) gives, by expanding the partial derivative,

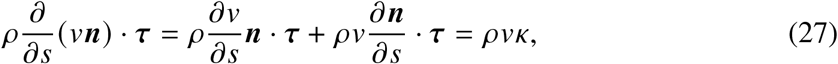

where the second equality holds because ***n*** and ***τ*** are perpendicular and *κ* = *∂* ***n***/*∂s* · ***τ*** is the signed curvature of the interface (Berger, 2003). Substituting 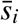 for an arbitrary, continuous arc-length coordinate *s*, and omitting arguments (*s, t*) from the notation, the evolution of spring density can be written as

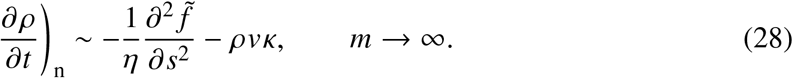

#### 2.2.1 Cell density

When the number of springs in the discrete model increases, spring den sity d iverges as O(*m*) and characteristic timescales of mechanical relaxation slow down as O(1/*m*^2^) (Buenzli et al., 2025). To obtain well-defined quantities and finite relaxation rates that are independent of *m* as *m* → ∞, some rescalings with *m* are required.

First, we define the local cell density *q* (*s, t*) at arc-length position *s* and time *t* such that

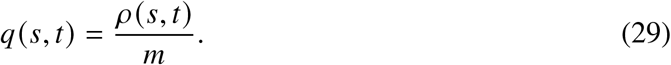

Substituting Equation (29) into Equation (28) gives

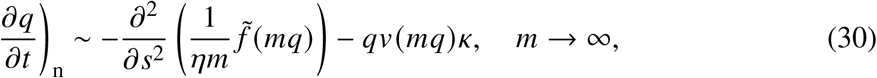

Where

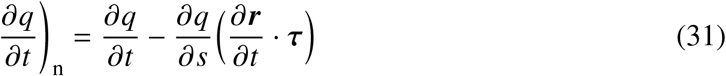

represents the rate of change of cell density following trajectories normal to the interface at all times. For Equation (30) to be well-defined in the limit *m* → ∞, the restoring force, drag coefficient and normal velocity must be such that the following limits exist:

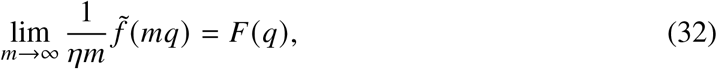

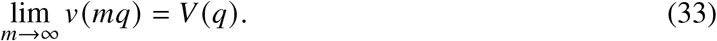

These limits define a cell-density-dependent restoring force *F* (*q*), as well as a cell-density-dependent normal velocity function *V* (*q*). The evolution of cell density, in the limit, is now given by

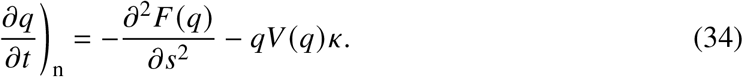

Equation (34) can be recast as a reaction–diffusion PDE with nonlinear diffusion,

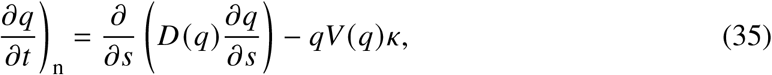

where the diffusive flux is related to the force gradient by

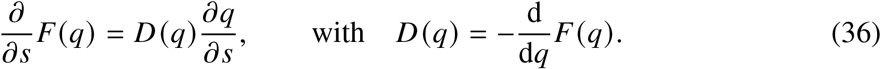

The diffusion term on the right hand side of Equation (35) represents changes in cell density as the tissue interface relaxes mechanically. The reaction term reveals that the evolution of cell density at the tissue scale is curvature-dependent due to cell-crowding effects during growth. This curvature dependence is an emergent property in the continuum limit, since curvature is not explicitly encoded in the discrete model at the cellular scale.

For the limits in Equations (32)–(33) to be well-defined, the parameters *k, η, a* and *k*_f_ from the discrete model in Equations (10)–(11) need to be rescaled with the number of springs per cell *m* (Buenzli et al., 2025). We now refer to these parameters with a superscript ‘(*m*)’ and rescale them according to

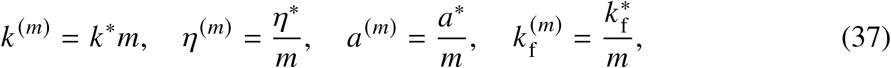

where *k*\*, *η*\* and *a*\* represent the mechanical properties of the entire cell, and 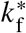 is the secretory rate of a single cell (area of new tissue secreted per unit time per cell). The scalings for *k* ^(*m*)^, *η*^(*m*)^, and *a*^(*m*)^ have been used previously in Murray et al. (2009) and Murphy et al. (2019) and rigorously justified in Buenzli et al. (2025). The scaling for 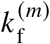 comes from substituting Equation (1) in Equation (33), which leads to

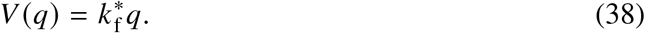

It represents the assumption that cells generate the same area of tissue material regardless of the number of springs within the cell (see Figure 2e).

With these parameter scalings and the Hookean restoring force in Equation (10), we have

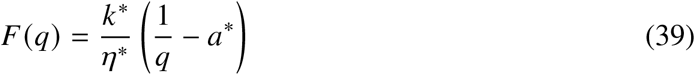

and a nonlinear diffusion with density-dependent diffusivity

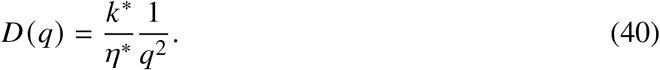

With the nonlinear restoring force in Equation (11), we have

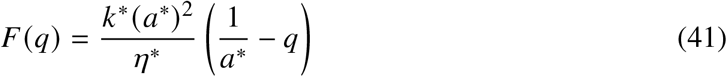

and a linear diffusion with constant diffusivity that is independent of density

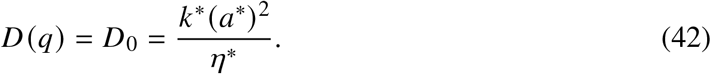

These diffusion models are consistent with previous works of mechanical relaxation of cells on static interfaces (Murray et al., 2009; Murphy et al., 2019; Buenzli et al., 2025). Both Equations (40) and (42) show that mechanical relaxation at the tissue scale depends on the mechanical properties of individual cells and is independent of geometric effects. When the nonlinear restoring force is used, the evolution equation of cell density in Equation (35) becomes

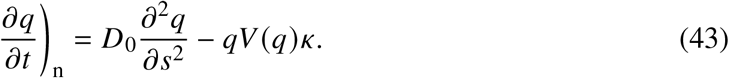

This equation has been derived in previous studies based on continuum models of cell crowding during tissue growth (Alias and Buenzli, 2017, 2019; Hegarty-Cremer et al., 2021). An advantage of our discrete model approach is to provide a cell mechanics justification for the ad-hoc addition of a diffusive term employed in these previous works, and the consideration of other diffusion laws based on single-cell mechanical properties.

### 2.3 Numerical simulations methods

We apply our models to simulate tissue growth on substrates of various shapes, including pores of square, hexagonal, and circular shapes, and trench-like geometries inspired by trabecular bone (Alias and Buenzli, 2017) and by tissue growth experiments of Bidan et al. (2012).

#### 2.3.1 Discrete model

Numerical solutions of the discrete model are obtained by solving the system of ordinary differential equations in Equation (2), where 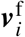 is given by either Equation (5) or Equation (6), and 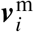 is given by Equation (9). These equations rely on a choice of a restoring force law *f* (*ℓ*) (Equation (10) or Equation (11)), as well as on a specification of the normal velocity *v*_*i*_, taken to be 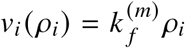 to correspond to the cell’s normal velocity *V* (*q*) in Equation (38). When the initial substrate interface is polygonal, we ensure that a spring boundary is placed initially at each vertex of the polygon. We then divide each edge of the polygon evenly to obtain *M* springs in total. To distribute spring boundaries evenly on substrate interfaces described by smooth curves, we divide arc-length reparametrisations calculated by numerical quadrature into *M* equidistant arc-length displacements (Buenzli et al., 2025).

After initialising spring node positions on the substrate interface, we solve the system of *M* ordinary differential Equations (2) using the `DifferentialEquations.jl` package (Rackauckas and Nie, 2017). Since Equation (3) depends on a constant time step Δ*t*, we solve the system using the Runge–Kutta 4 method with the `RK4` algorithm and a constant time step of Δ*t* = 0.01 days. In practice, we calculate 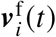 algorithmically by first determining the position of the new node 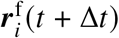 after a time increment Δ*t*, and then determining 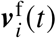 using Equation (3). The open-source *Julia* code that implements this numerical scheme is available at Kuba et al. (2026).

#### 2.3.2 Continuum model

Numerical simulations of the continuum model are conducted for the nonlinear restoring force leading to the evolution of cell density in Equation (43) with constant diffusivity, coupled with the interface’s normal velocity in Equation (1). Since the tissue interface is a closed curve, periodic periodic boundary conditions are employed. To track the evolution of the tissue interface *S*(*t*), we use a time-dependent polar curve parametrization

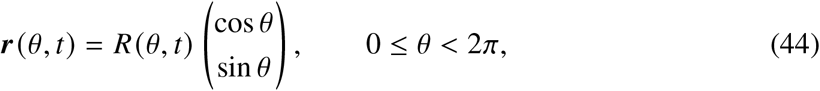

with *R*(0, *t*) = *R*(2*π, t*). The interface evolves according to the normal velocity condition together with the cell density equation,

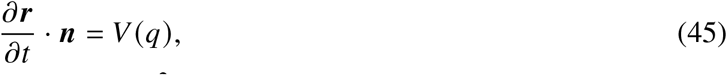

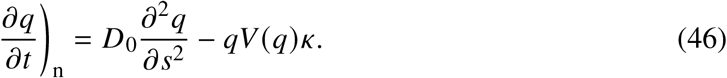

Equations (45)–(46) are solved numerically using the schemes devised by Alias and Buenzli (2017), re-implemented in *Julia*. At intermediate and high diffusivities, Equation (46) is solved using a semi-implicit finite difference scheme where the curvature-dependent term is solved explicitly (forward Euler) and the diffusion term is solved implicitly (backward Euler). At low diffusivities, Equation (46) is solved in conservative form using the finite volume method with semi-discrete Kurganov–Tadmor scheme (Kurganov and Tadmor, 2000) and a fully explicit forward Euler discretisation in time. A more detailed description of the discretisation and numerical scheme is presented in Appendix B. The open-source *Julia* code that implements these numerical methods is available at Kuba et al. (2026).

### 2.4 Parameter values

To obtain biologically realistic simulations, we estimate the value of 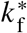 by referring to the experimental results of *in vitro* tissue growth within square pores with a side length of 500 μm from Buenzli et al. (2020). These pores have an initial area of 2.5 × 10^5^ μm^2^, and were found to close due to tissue growth after *T*_b_ ≈ 28.46 days, called the bridging time.

Since each cell in our model is assumed to produce a constant area of new tissue 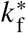 per unit time, the total amount of tissue produced by time *t* by all *N* cells in our model is given by

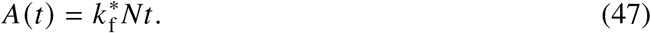

The remaining void area in the pore, *Ω*(*t*) is therefore given by

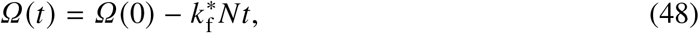

where *Ω*(0) is the initial pore area. We further set *N* = 100 by assuming that cells have an initial length of 20 μm (Simpson et al., 2010, 2013; Qiu et al., 2019; Lanaro et al., 2021) and that they form a confluent layer on the perimeter of the 500 μm × 500 μm square pore. The parameter 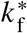 is then chosen such that *Ω*(*T*_b_) = 0, giving 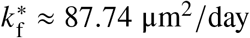. We use this value of 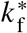 for all numerical simulations.

In the discrete model, the rate of mechanical relaxation is proportional to the ratio *k*/*η* (Buenzli et al., 2025). For all simulations, we choose *η* = 1, and explore how variations of spring stiffness *k* and resting length *a* affect the evolution of the tissue interface in Section 3. These parameter values influence the diffusivity in the continuum model as per Equations (40) and (42).

## 3 Results & Discussion

We start by numerically simulating tissue growth in a square pore of size 100 μm× 100 μm using both the discrete and continuum models with the nonlinear restoring force. We choose an initial cell density of *q*_0_ = 0.05 μm^−1^, corresponding to distributing *N* = 20 cells of size 20 μm evenly along the square interface. Figure 3a shows numerical simulations of the discrete model when the cells contain either a single spring (*m* = 1) or multiple springs per cell (*m* = 4). The initial perimeter of the square in these simulations is chosen to be small to limit the total number of cells and allowing us to clearly visualise cell boundaries (black crosses) and spring boundaries (grey dots) along the initial interface. The interface of the tissue is shown at regular time intervals as solid curves coloured by the local cell density. Cell positions evolve as they deposit tissue and interact mechanically, leading to the rounding behaviour of the tissue interface observed experimentally (Figure 3a). Rounding is due to crowding effects that increase cell density where curvature is large near the corners of the square, and to the fact that the normal velocity of the interface is proportional to cell density, see Equation (38).

**Figure 3.**
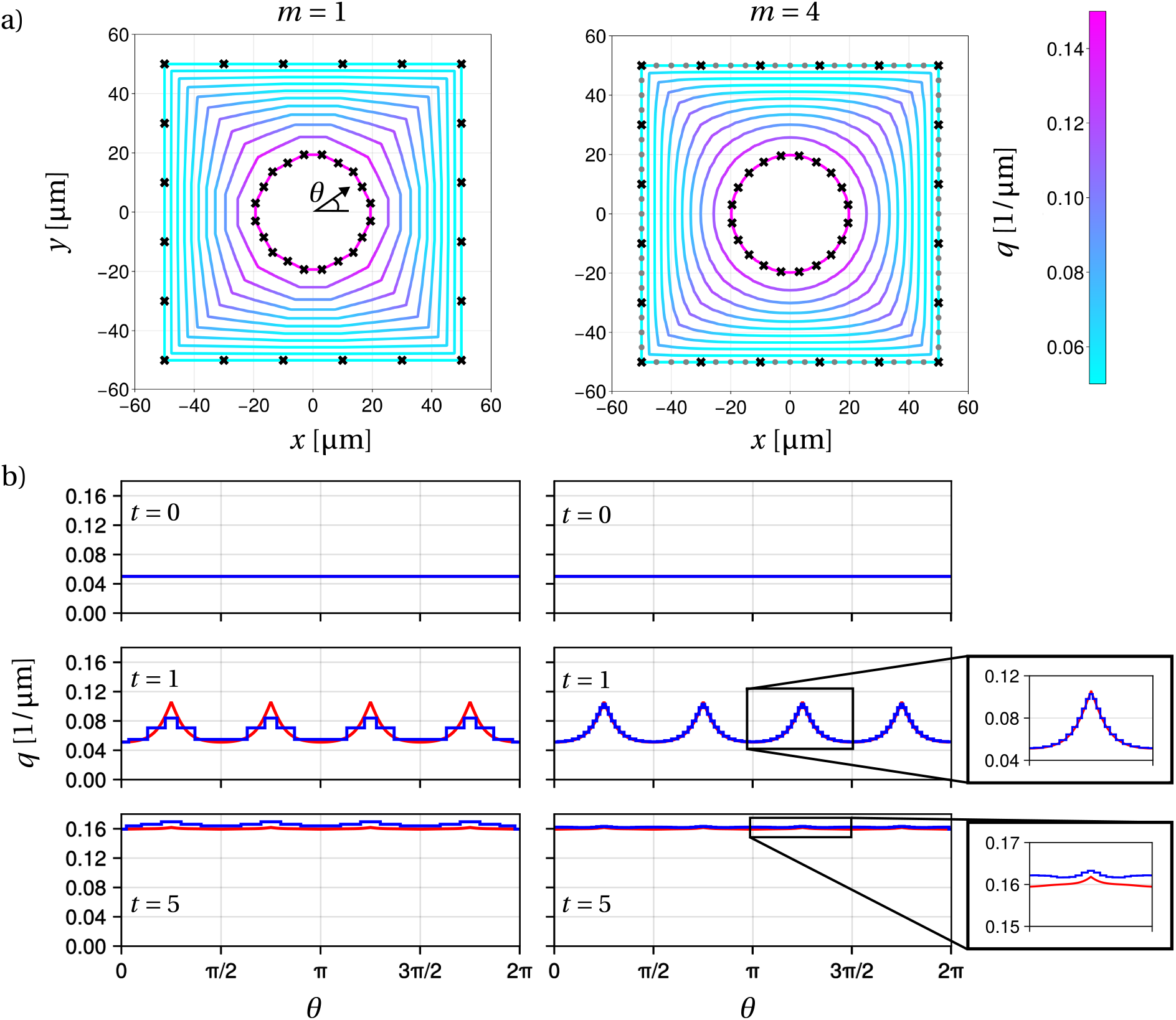
Simulations of the discrete model with *N* = 20 cells. (a) Each solid line represents the tissue interface at regular time intervals of 0.5 days coloured by the discrete cell density during a 5 day simulation period, demonstrating interface smoothing. Discrete cell boundaries are shown at *t* = 0 with black crosses. In the case of *m* = 4 springs per cell, spring boundaries are shown at *t* = 0 as grey dots. (b) Discrete (blue) and continuous (red) cell density profiles along the angular position at different simulation times. The initial cell density is *q*_0_ = 0.05 μm^−1^. The mechanical relaxation rate corresponds to a diffusivity *D* = 100 μm^2^/day via Equation (42) with *k*\* = 100 *N*/μm, *η*\* = 1 *N* day/μm, and *a*\* = 1 μm. Each cell produces 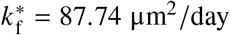.

While the final position of cell boundaries is not affected significantly by the value of *m*, the interface and cell density heterogeneities becomes smoother with larger *m* (Figure 3a). Figure 3b compares cell density profiles at angle *θ* arising from the discrete model (blue) with cell density profiles arising from the continuum model (red), at times *t* = 0, 1, 5 days. While simulations with *m* = 1 adequately approximate the numerical solution of the continuum model, simulations with *m* = 4 reduce discrepancies between discrete and continuous cell densities.

A comparison of the evolution of the tissue interface between the discrete and continuum models is provided in Figure 4, where we explore the effects of varying the mechanical relaxation rate *k*\*/*η*\* and corresponding diffusivity *D* = *k*\*(*a*\*)^2^/*η*\* from Equation (42). To provide an accurate match between the discrete and continuum models, we set *m* = 10. We also set the cell resting length to be *a*\* = 10 μm, so that cells are in tension initially (Bidan et al., 2016; Lanaro et al., 2021). Figure 4 shows simulations performed with *D* = *k*\*(*a*\*)^2^/*η*\* = 1 μm^2^/day (slow relaxation rate, small diffusivity), 100 μm^2^/day (intermediate relaxation rate and diffusivity), and 10^5^ μm^2^/day (fast relaxation rate, large diffusivity).

**Figure 4.**
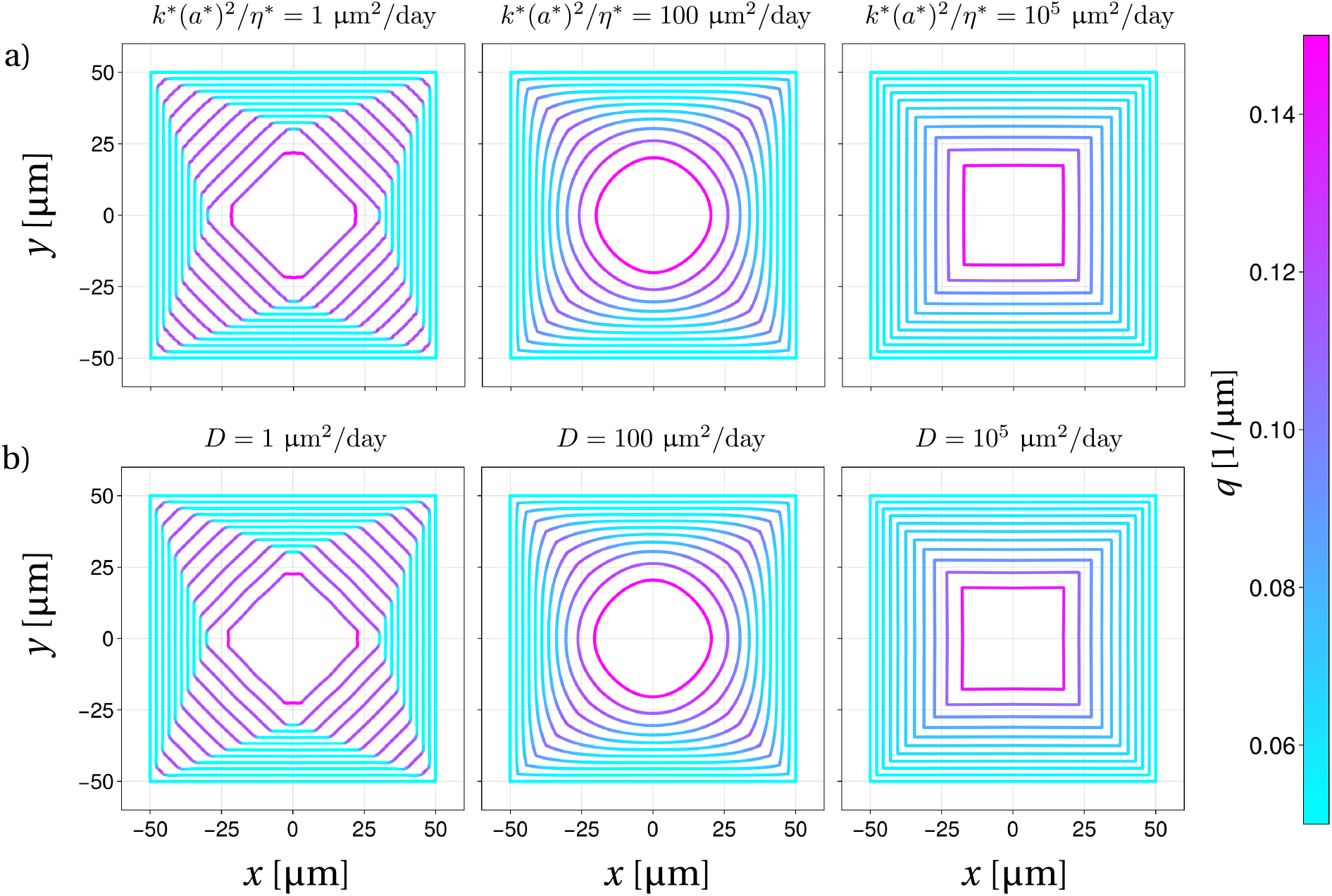
Simulations of the discrete (a) and continuum (b) models with low, intermediate, and high values of *k*\* (*a*\*)^2^ *η*\* and *D*, respectively. Simulations are conducted across low (*D* = 1 μm^2^ day), intermediate (*D* = 100 μm^2^ day), and high (*D* = 10^5^ μm^2^ day) diffusivities for the continuum model and their corresponding values of *k*\* (*a*\*)^2^ *η*\* given by Equation (42). The interface is recorded at regular intervals of *Δt* = 0.5 days during a 5 day simulation period with colour indicating cell density. For the discrete model, simulations are performed using *N* = 20 cells each comprised of *m* = 10 springs, with an initial cell density *q*_0_ = 0.05 1/μm. Cells have a resting length *a*\* = 10 μm, and produce an area of tissue at a rate of 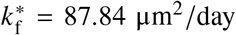. Diffusivity coefficients are set by varying the cell stiffness *k*\* and holding *η*\* = 1 *N* day μm as a constant.

The discrete model simulations (Figure 4a) and continuum model simulations (Figure 4b) produce visually identical results across all values of *k*\*(*a*\*)^2^/*η*\* considered. Slow mechanical relaxation and small diffusivity (*k*\*(*a*\*)^2^/*η*\* = 1 μm^2^/day) leads to cells rapidly accumulating near sharp corners. This creates high-density regions and cusps in the interface that propagate laterally in the form of shock waves due to differences in interface velocities (Alias, 2017). These cusps smooth out with intermediate values of *k*\*(*a*\*)^2^/*η*\* = 100 μm^2^/day, resulting in a rounded tissue interface consistent with experimental observations in *in-vitro* cell culture experiments and in bone tissue formation (Rumpler et al., 2008; Bidan et al., 2012, 2013; Alias and Buenzli, 2017), as seen in Figure 1. Fast mechanical relaxation rates and large diffusivity (*k*\*(*a*\*)^2^/*η*\* = 10^5^ μm^2^/day) preserve cusps of the initial substrate since cell density remains homogeneous throughout growth and the tissue interface evolves by uniform offsets. It is interesting to observe that while numerical simulations of the continuum model require sophisticated finite-volume numerical techniques at low diffusivity, discrete model simulations can use a standard ODE solver for any value of *k*\*(*a*\*)^2^/*η*\*. Since the discrete model converges to the continuum model as *m* → ∞ (Section 2.2), the discrete model can be used as a novel discretisation method of the continuum reaction–diffusion model in Equation (35).

A major advantage of the discrete model is its ability to track the positions and trajectories of cells for any restoring force and interface geometry. This individual-level detail cannot be provided by standard implementations of the continuum model. In Figure 5, we show the trajectories of a selected number of cells during new tissue growth for three different substrate geometries (circle, square, and trench-like). We compare simulations using either the Hookean restoring force in Equation (10) or the nonlinear restoring force in Equation (11). In all simulations, we take a uniform initial cell density of *q*_0_ = 0.05 μm^−1^. We choose a value of *k*\*/*η*\* = 10 μm^2^/day, such that for both restoring forces, the diffusivities in Equation (40) and (42) are equal at time *t* = 0 and correspond to the intermediate diffusivity regime described in Figure 4. To simplify visualisation of cell and spring trajectories, simulations were conducted with *m* = 2. Black lines represent the trajectories of cell boundaries and red lines representing the trajectory of the internal spring boundary. In the circle and square pores, we observe that despite minor variations in the trajectories of cell and spring boundaries, the overall rounding behaviour of the tissue interface remains similar between the two restoring forces. In the trench-like geometry, there are more noticeable variations in the trajectories of cell and spring boundaries at early and intermediate times.

**Figure 5.**
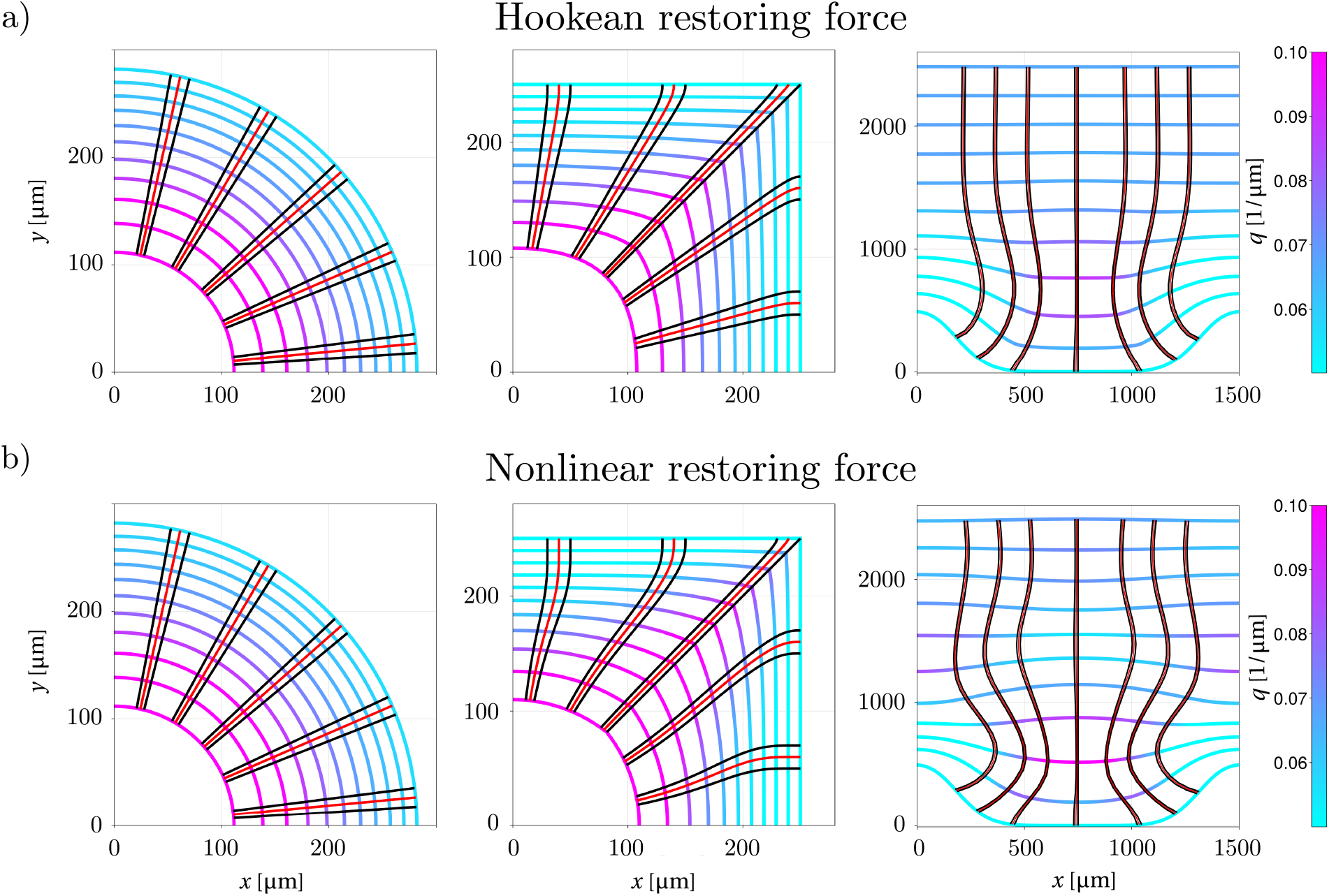
Numerical simulations of the discrete model for square, circular and trench-like geometries using the Hookean restoring force (a) and nonlinear restoring force (b). The tissue interface is shown at regular time intervals Δ*T* = 2.4 days for the square and circular geometries, and Δ*T* = 40 days for the trench-like geometries (coloured curves). A selected number of cell boundary trajectories are shown (solid black lines) along with their inner spring boundary (solid red lines). Simulations are performed using *N* = 100 cells each comprised of *m* = 2 springs, with an initial cell density *q*_0_ = 0.05 1/μm. Cells have a resting length *a*\* = 10 μm, and produce an area of tissue at a rate of 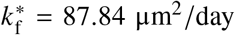. To ensure interface rounding in confining geometries the mechanical relaxation rate for Hookean springs corresponds to an initial diffusivity *D* (*q*_0_)= 10 μm^2^ day via Equation (40), and *D*_0_ = 100 μm^2^ day for nonlinear springs via Equation (42).

The mechanical stress state of a cell in the simulations depends significantly on the cell resting length *a*\* assumed. In Figure 6, we compare simulations of the discrete model with Hookean force (Figure 6a) and with nonlinear force (Figure 6b) for various values of *a*\*. We choose values of *a*\* such that cells are initially either in tension (*a*\* = 10 μm), stress free (*a*\* = 20 μm), or in compression (*a*\* = 30 μm). To enable a consistent comparison across different values of *a*\*, we ensure that the initial diffusivity is identical for each restoring force. For Hookean springs, the diffusivity in Equation (40) is independent of *a*\*, so that we use the same value of *k*\*/*η*\* for all values of *a*\*. In contrast, for the nonlinear spring model, diffusivity depends on *a*\* according to Equation (42), so we set *k*\*/*η*\* = *D*_0_/(*a*\*)^2^. With these choices of restoring force laws and parameters, despite mechanics and geometry being coupled during tissue growth, identical tissue evolutions may be obtained with different cellular stress states. These results indicate that cell resting length does not affect the evolution of the tissue interface in the continuum limit as *m* → ∞. Contributions due to the resting length cancel in the force balance at a spring boundary in Equation (18). However, depending on the value of *a*\*, cells may find themselves under tension (green) or compression (blue).

**Figure 6.**
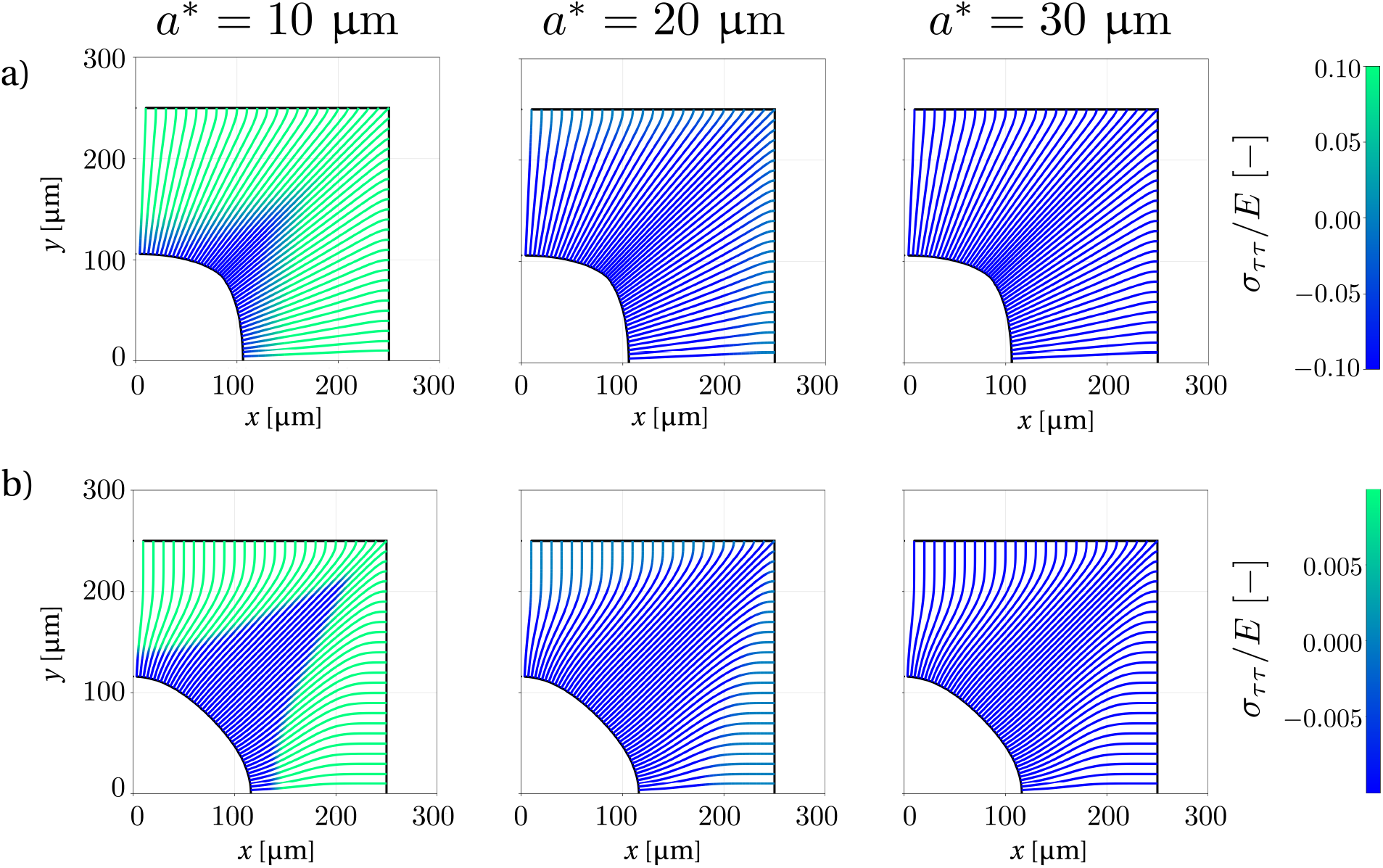
Cell boundary trajectories of the discrete model coloured by normalised stress *σ*_*ττ*_ *E* defined in Equation (12). Only the top-right quarter of the domain is shown. Numerical simulations are performed for different cell resting lengths: *a*\* = 10 μm (cells initially under tension), *a*\* = 20 μm (cells initially stress-free), and *a*\* = 30 μm (cells initially under compression). The tissue interfaces at *t* = 0 and *t* = 24 days are shown as solid black lines and show that identical tissue evolution are obtained despite cells experiencing different mechanical stresses. For each of the restoring forces, *k*\* is scaled using Equations (42) and (40) such that for the Hookean springs, *D* (*q*_0_)= 10 μm^2^ day, and for nonlinear springs, *D*_0_ = 100 μm^2^ day. Simulations are performed using *N* = 100 cells each comprised of *m* = 10 springs, with an initial cell density *q*_0_ = 0.05 1/μm, and a tissue production rate 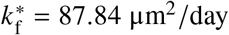.

An important consideration in tissue engineering constructs is the time required for new tissue to grow into porous scaffolds. In a previous study of tissue growth in 3D-printed porous scaffolds with square pores of size *L*, we showed that the so-called bridging time *T*_b_ needed for new biological tissue to fill the square pores scaled linearly with *L*, suggesting that individual cell behaviours such as cell proliferation and cell migration were unaffected by pore size (Buenzli et al., 2020). The mathematical model presented in our present work allows us to refine the relationship between bridging time *T*_b_ and pore size by accounting for the influence of pore shape. In our discrete model, cells are assumed to produce tissue at a constant rate 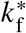, meaning that a given number of cells generate the same area of tissue regardless of their initial arrangement. The initial number of cells is *N* (*L*) = *q*_0_ *P*(*L*), where *q*_0_ is the initial cell density, and *P*(*L*) is the perimeter of the initial scaffold pore of linear size *L*. By substituting this expression for *N* (*L*) into Equation (48) and rewriting the initial pore area as *Ω*(0) = *A*(*L*) to emphasise its dependence upon pore size *L*, we obtain

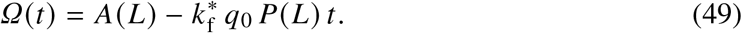

Since by definition of bridging time, *Ω*(*T*_b_) = 0, we obtain from Equation (49) a relationship showing that the size dependence of *T*_b_ originates from the ratio of initial pore area and initial pore perimeter:

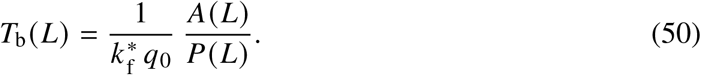

Equation (50) provides a prediction of bridging time for arbitrary 2D pore shapes, which is a significant generalisation of our previous finding that bridging time scales linearly with pore size in square pores (Buenzli et al., 2020). While the ratio of pore area over pore perimeter in Equation (50) always scales linearly with *L* for regular (nonfractal) shapes, the scaling is modulated by the shape factor *A*(*L*)/*P*(*L*). For instance, for square pores of size *L, A*(*L*) = *L*^2^ and *P*(*L*) = 4*L*, so that

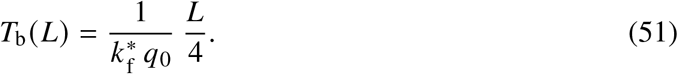

However, for hexagonal pores of maximal diameter 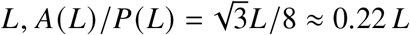. Because our model does not include mechanobiological regulation of cellular activities, the influence of pore shape for bridging time provided by Equation (50) is entirely attributed to geometric crowding effects.

In Figure 7 we compare the void area *Ω*(*t*) of square pores with side lengths *L* = 500, 750, 1000 μm and initial cell density *q*_0_ = 0.05 μm^−1^ with the analytic expression given by Equation (49). The void area enclosed by the discrete tissue interface, is estimated using the *shoelace formula* (Lee and Lim, 2017). The results show good agreement between the discrete model estimates and the analytic solution. While a larger initial pore means a larger void area to fill, the rate of closure is faster as it is directly proportional to the initial perimeter *P*(*L*), see Equation (49).

**Figure 7.**
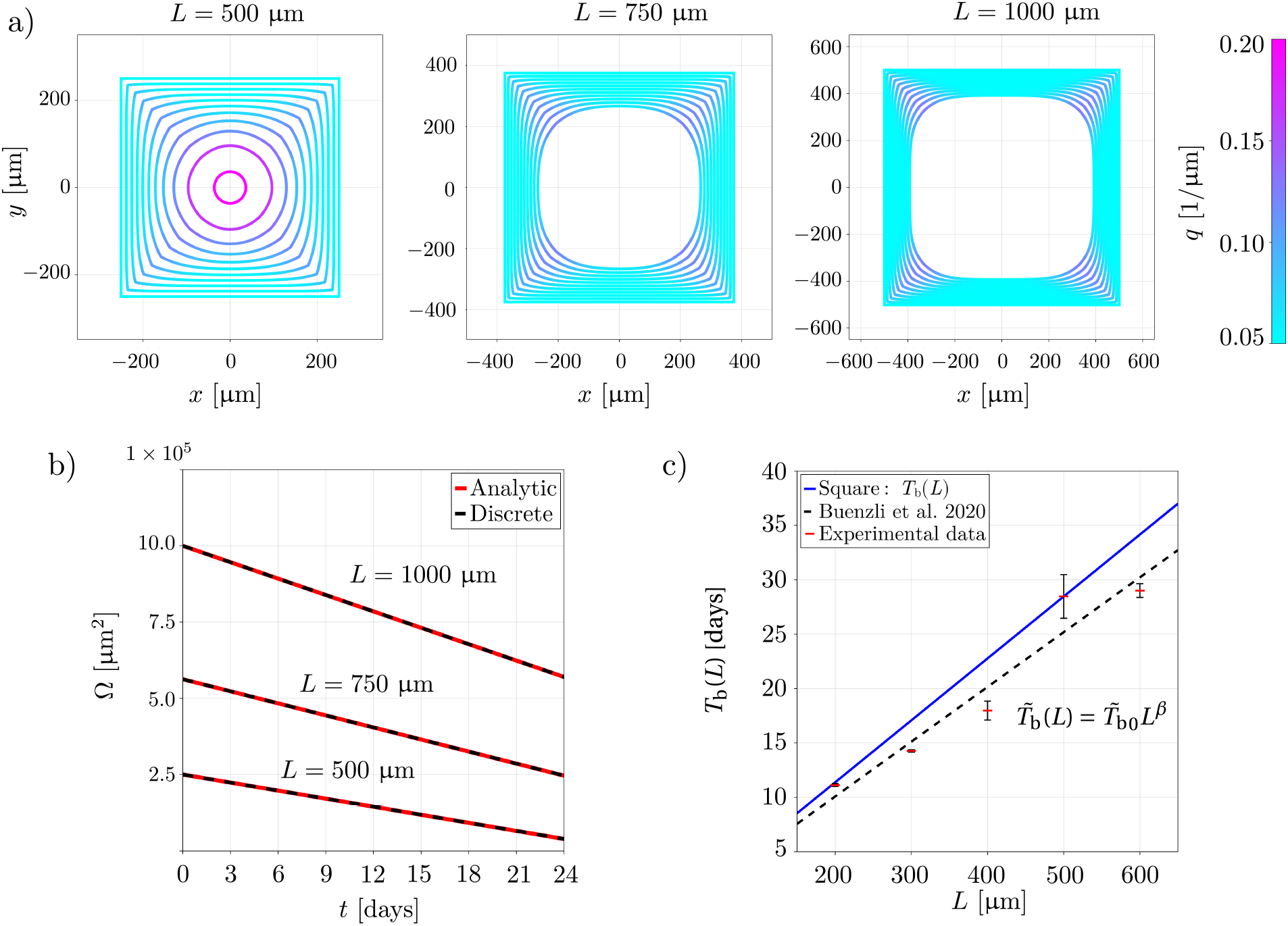
Simulations of tissue growth within a square pore geometry with different initial side lengths *L*. (a) Tissue interface at regular time intervals of 0.5 days. All simulations use the same initial cell density of *q*_0_ = 0.05 μm^−1^. (b) Void area *Ω* (*t*), comparing simulation results (solid black line) and the analytic expression in Equation (49) (dashed red line). (c) Bridging time as a function of side length derived from the geometric properties of a square pore (blue), compared with the regression model from Buenzli et al. (2020) (black), calibrated on experimental data (red). The regression model parameters are 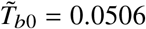 and *β* = 0.9990. Error bars correspond to the standard error on *T*_b_ provided by fitting experimental time evolutions of *Ω*(*t*) to the function 1 − (*t*/*T*_b_)^*ν*^ (Buenzli et al., 2020).

Both the regression model of Buenzli et al. (2020) and Equation (50) predict a bridging time proportional to *L* (Figure 7c). While the regression model is an empirical power-law fit to experimental data, the linear dependence in Equation (50) emerges directly from our discrete model, providing support for its underlying assumptions. Our model slightly overestimates the bridging time in Figure 7c because the tissue formation rate *k*_f_ was calibrated using only the 500 μm × 500 μm pore data. In contrast, the regression model of Buenzli et al. (2020) was fitted to data from all pore sizes.

## 4 Conclusions

The observation that biological tissues grow at rates that can be influenced by the local geometry of the tissue interface has spurred high interest in tissue engineering and developmental biology in the last two decades (Callens et al., 2020; Schamberger et al., 2023). The influence of curvature in tissue growth has been suggested to arise due to geometric crowding effects due to tissue growing into restricted available space (Alias and Buenzli, 2017, 2018, 2019; Buenzli et al., 2020; Hegarty-Cremer et al., 2021), mechanobiological effects such as increased proliferation of cells experiencing higher mechanical stress due to their geometric configurations (Nelson et al., 2005; Dunlop et al., 2010; Lim et al., 2010; Nelson, 2022), as well as surface tension effects (Rumpler et al., 2008; Bidan et al., 2012, 2013; Ladoux and Mège, 2017; Ehrig et al., 2019). Most previous modelling approaches use continuum frameworks to explore the relationship between tissue growth rate and geometry or mechanics. These models can be challenging to connect with experimental measurements of cell positions or their trajectories, since these cell-level details are not modelled in continuum frameworks. Our work introduces a novel, comprehensive mathematical model of discrete cells to simulate the evolution of tissue interfaces during growth. The proposed model significantly extends previous discrete models of mechanical cell interactions by coupling curvature-induced crowding effects and mechanical relaxation, while allowing mathematically tractable analyses. While the present paper focuses on developing the theoretical and computational framework of this model, future works using this framework will be able to connect more directly with cell-level experimental data.

A major result of our work is the analytical derivation of curvature-dependent tissue growth from a discrete model of cells interacting mechanically. This derivation relies on identifying appropriate microscopic rules governing cell–cell interactions during tissue growth that accurately represent the crowding generated by new tissue production. These rules depend on the relative orientation of neighbouring cells. Alternative evolution rules do not in general recover the physically relevant viscous (or entropic) solution of Equations (45)–(46) (Sethian, 1999; Alias and Buenzli, 2017). In the continuum limit, cell density satisfies a reaction–diffusion equation that recovers the continuum cell population model of Alias and Buenzli (2017), while providing a mechanistic derivation of its diffusion term. More broadly, our framework provides a direct link between individual cell properties, such as mechanical stiffness and tissue production rate, and emergent tissue-scale behaviour, including the diffusive tissue mechanical relaxation rate and curvature-dependent evolution. Numerical simulations of our model show that growth and mechanics interact strongly and that changing their relative rates leads to widely different tissue evolutions. Finally, our model provides a theoretical prediction for how fast pores of arbitrary shapes in tissue engineering scaffolds can be bridged by new tissue growth based on the ratio of the initial pore area and initial pore perimeter. This result refines previous experimental findings that pores require a bridging time that scales linearly with size (Buenzli et al., 2020), by accounting for a pore shape factor.

### 4.1 Model limitations

Our model has a number of limitations:

- The tissue is assumed to be two-dimensional, with the tissue interface represented by a curve. This corresponds to modelling a tissue in cross-section, and is appropriate to model some cell cultures in thin 3D-printed scaffolds, where cells grow thin tissue layers, and where edge effects do not introduce significant three-dimensional behaviour. However, the model cannot account for more complex three-dimensional tissue geometries involving out-of-plane curvatures (Ehrig et al., 2019; Callens et al., 2020; Schamberger et al., 2023).
- New tissue material is assumed to be produced near the tissue interface, representing surface growth. Tissue production is specified as an area of new tissue produced per cell at the interface, which displaces the cells in the normal direction at a rate proportional to cell density (Buenzli, 2015). The model therefore does not currently describe bulk growth, in which new tissue material is produced within deeper tissue layers.
- Cell mechanics is represented by simple, spring-like mechanical interactions between adjacent cell boundaries. The mechanical response of real cells is much more complex and involves multiple biophysical processes including actin polymerisation, membrane adhesion complexes, and cytoplasmic hydrostatic pressure (Moeendarbary and Harris, 2014; Ladoux and Mège, 2017), none of which are explicitly represented in the model. However, a spring-based model provides a simple description of mechanical interactions between neighbouring cells, capturing both force transmission and cell body deformation when multiple springs per cell are considered.
- Finally, the model does not explicitly represent cell proliferation, death, or differentiation. Instead, the rate of new tissue production per cell, 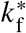, represents newly formed tissue material. This material may implicitly represent both extracellular matrix and cells that arise through proliferation and differentiation and subsequently become embedded within the tissue.

### 4.2 Future works

The present work establishes a theoretical and computational framework for describing how cell activities and mechanical relaxation interact during tissue growth. There are many opportunities to extend this work and to strengthen its relationship with biological experiments. The discrete framework proposed provides new opportunities to link tissue level experimental data, such as rates of mechanical relaxation, with individual cell-level properties such as cell stiffness, that may be challenging to measure directly. For example, by describing individual cells explicitly, our model opens up the possibility of using live imaging and cell tracking technologies as potential avenues to infer effective mechanical properties of the cells during tissue growth, such as their intrinsic restoring force law. In its current formulation, the discrete model assumes that the tissue interface is represented by a constant number of cells, representing a balance between cell proliferation, and cell embedment into new tissue material. Incorporating cell proliferation, differentiation, and death more explicitly into the model would allow us to modulate the number of cells that make up the tissue forming interface and to model the composition of the tissue generated. This will help connect bulk properties of the tissue, such as tissue-embedded cell density, with experimental data extracted from tissue growth experiments or from biological tissue samples using parameter inference techniques (Kreutz et al., 2013; Alon, 2019; VandenHeuvel et al., 2023; Murphy et al., 2024; Simpson and Baker, 2026). Further extensions may also include the consideration of individual cell shapes and orientations, to connect more directly with data collected in cell culturing experiments and bone histology studies (Marotti et al., 1975; Bartnikowski et al., 2014; Lanaro et al., 2021; Devlin et al., 2024). These extensions will be the subject of future works.

## Acknowledgements

This research was supported by the Australian Research Council (DP190102545, DP230100025). SK and PRB acknowledge support from the Max Planck Queensland Centre for the Materials Science of Extracellular Matrices. We thank Brenna Devlin and Maria A. Woodruff for providing the experimental image presented in Figure 1a. We also thank Almie M. Alias for the original implementation in MATLAB of the numerical algorithm described in (Alias and Buenzli, 2017).

## Appendix A Continuum limit

In this appendix we examine the asymptotic behaviour of Equation (20) as *m* → ∞. With Equations (24)–(25), we can rewrite Equation (20) as

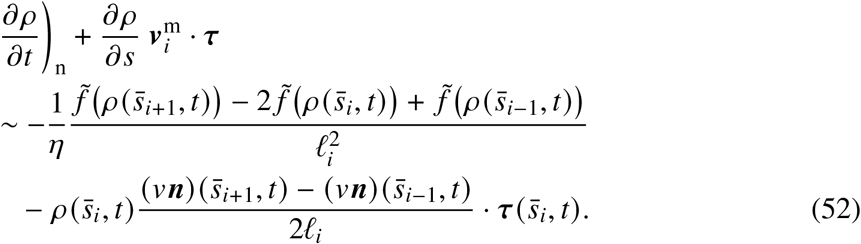

Buenzli et al. (2025) showed that the second term in the left hand side of Equation (52) cancels out precisely with some terms arising from expanding the first term in the right hand side of Equation (52) about 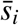. As in Buenzli et al. (2025), we introduce a parametrisa tion 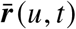 of the interface that tracks the spring midpoints at all times, such that 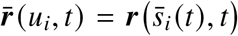 where *u*_*i*_ = *i*Δ*u, i* = 1, …, *M*, are regularly space and time independent. The arc-length coordinates of the spring midpoints are given by

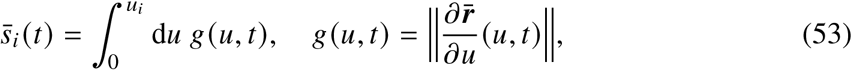

so that

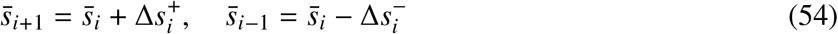

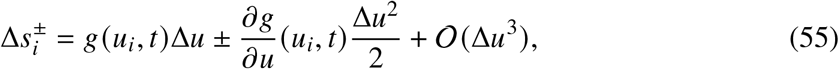

From Equation (53), spring density at arc-length coordinate 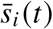, which corresponds to coordinate *u*_*i*_, is given by,

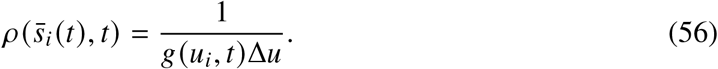

Using Equation (53) and (56), we have

(*∂ ρ*/*∂s*) = −(*∂g*/*∂s*)/(*g*^2^Δ*u*) = −(*∂g*/*∂u*)/(*g*^3^Δ*u*). Expanding 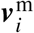.***τ*** about *u*_*i*_ using Equation (18), gives 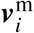.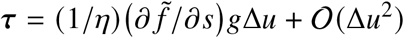, so that

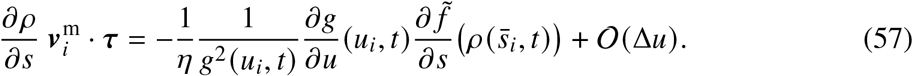

The expansion of the first line in the right hand side of Equation (52) about *u*_*i*_ as Δ*u* → 0 (*m* → ∞) gives (Buenzli et al., 2025):

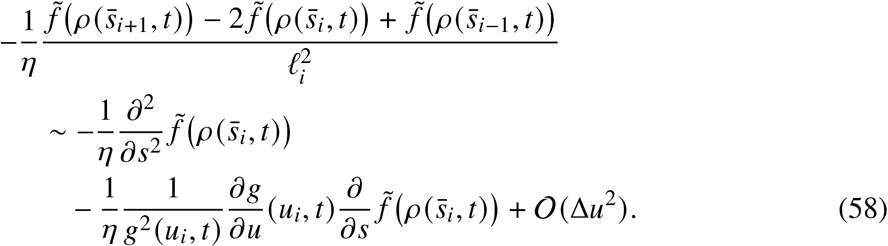

The second term in the right hand side of Equation (58) is identical to Equation (57), so that substituting Equations (57)–(58) into Equation (52) gives

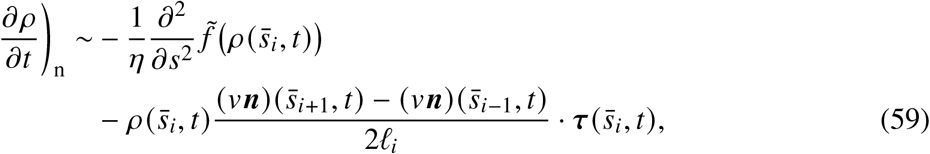

as *m* → ∞. We now examine the asymptotic behaviour of the second line in the right hand side of Equation (59) as *m* → ∞, which corresponds to changes in local density due to new tissue formation. Expanding this contribution about 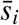 as Δ*u* → 0 using Equation (55) gives

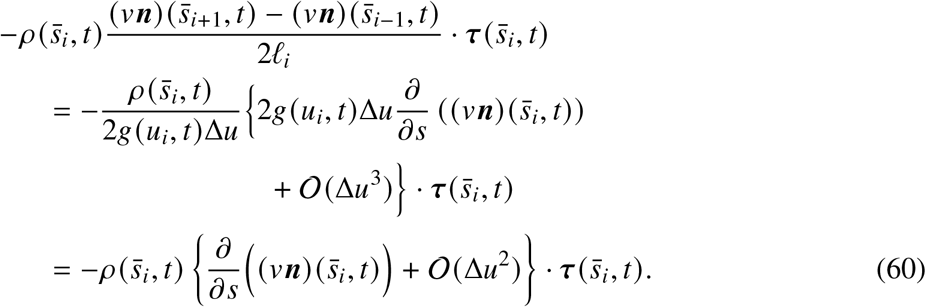

Therefore, Equation (59) becomes

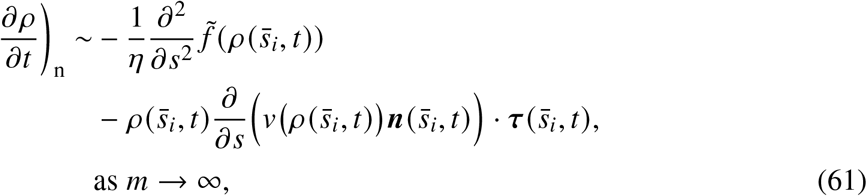

which is the same as Equation (26).

## Appendix B Continuum model simulations

We provide additional detail on the numerical scheme used in (Alias and Buenzli, 2017) to solve Equations (45)–(46) at intermediate–high and low diffusivities. To simplify notation, we write subscripts for partial derivatives, e.g., *q*_*θ*_ = *∂q*/*∂θ*. Using the polar parameterisation ***r*** (*θ, t*) in Section 2.3.2, Equations (45)–(46) become

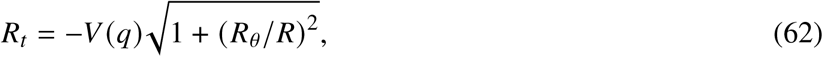

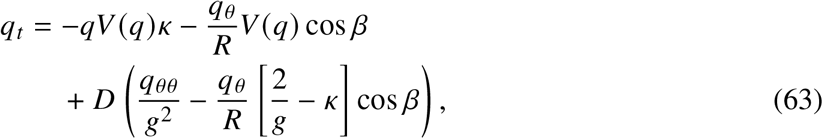

Where 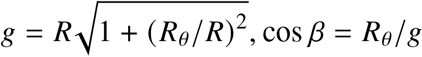, and 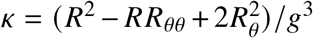. Equations (62)–(63) are solved using a semi-implicit finite difference scheme, with advective and reaction terms solved explicitly using forward Euler discretisation and diffusive terms solved implicitly using backward discretisation to limit restrictive Courant–Friedrichs–Lewy (CFL) conditions at high diffusivities (LeVeque, 2002). Central differencing is applied to all second-order derivatives in *θ*. Upwinding is applied to all first-order derivatives in *θ* according to the local sign of *R*_*θ*_, i.e.,

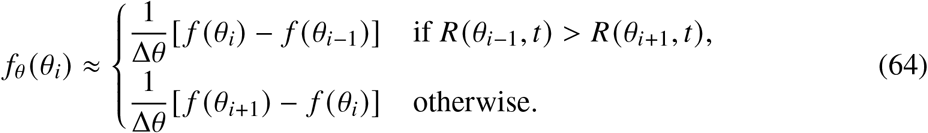

At low diffusivities, the accuracy of the simple finite difference scheme above deteriorates and a conservative scheme is required. Defining *σ* = *R*_*θ*_ and *ζ* = *qg*, Equation (46) is rewritten in conservative form, leading to the system of equations

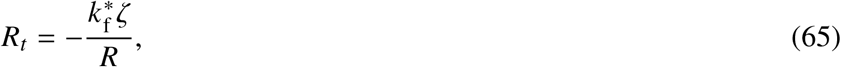

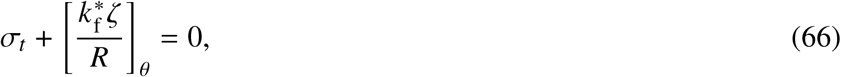

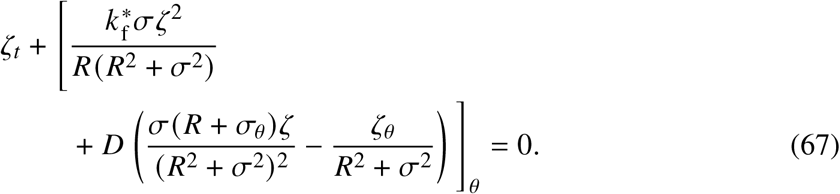

Equations (65)–(67) are recast in the form

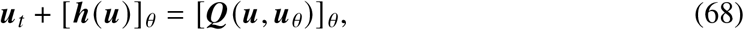

where ***u*** = (*R, σ, ζ*), ***h*** is hyperbolic and contains the part of the total flux that is independent of *ζ*_*θ*_, and ***Q*** is parabolic and contains the part of the flux in Equation (67) that depends on *ζ*_*θ*_. Equation (68) is discretised using the semi-discrete Kurganov-Tadmor scheme (Kurganov and Tadmor, 2000):

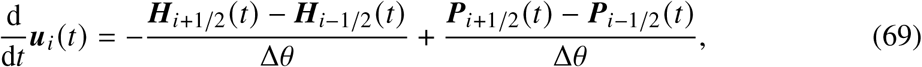

where ***H*** is the Rusanov numerical flux, and ***P*** is a second-order central difference approximation to the parabolic flux. Left and right interface values ***u***_*i*±1/2_ are interpolated via a minmod limiter, with the maximum eigenvalue of ***h***^′^(***u***) determined numerically. Finally, time integration of Equation (69) is performed using the forward Euler scheme. For a more detailed description of rewriting Equation (46) in conservative form and the implementation of the semi-discrete Kurganov-Tadmor scheme, we refer the reader to Alias (2017).

## References

Alias, M.A., 2017. Mathematical Modelling of the Geometric Control of Bone Tissue Growth. Ph.D. thesis. School of Mathematical Sciences, Monash University, Australia.

Alias, M.A., Buenzli, P.R., 2017. Modeling the effect of curvature on the collective behavior of cells growing new tissue. Biophysical Journal 112, 193–204. doi:10.1016/j.bpj.2016.11.3203.

Alias, M.A., Buenzli, P.R., 2018. Osteoblasts infill irregular pores under curvature and porosity controls. Biomechanics and Modeling in Mechanobiology 17, 1357–1371. doi:10.1007/s10237-018-1031-x.

Alias, M.A., Buenzli, P.R., 2019. A level-set method for the evolution of cells and tissue during curvature-controlled growth. International Journal of Numerical Methods in Biomedical Engineering 36. doi:10.1002/cnm.3279.

Alon, U., 2019. An Introduction to Systems Biology: Design Principles of Biological Circuits. 2 1ed., Chapman and Hall/CRC, New York.

Ambrosi, D., Ben Amar, M., Cyron, C.J., DeSimone, A., Goriely, A., Humphrey, J.D., Kuhl, E., 2019. Growth and remodelling of living tissues: perspectives, challenges and opportunities. Journal of The Royal Society Interface 16, 20190233. doi:10.1098/rsif.2019.0233.

Baker, R.E., Parker, A., Simpson, M.J., 2019. A free boundary model of epithelial dynamics. Journal of Theoretical Biology 481, 61–74. doi:10.1016/j.jtbi.2018.12.025.

Bartnikowski, M., Klein, T.J., Melchels, F.P., Woodruff, M.A., 2014. Effects of scaffold architecture on mechanical characteristics and osteoblast response to static and perfusion bioreactor cultures. Biotechnology and Bioengineering 111, 1440–1451. doi:10.1002/bit.25200.

Berger, M., 2003. A Panoramic View of Riemannian Geometry. Springer, Berlin, Heidelberg.

Bi, D., Lopez, J.H., Schwarz, J.M., Manning, M.L., 2015. A density-independent rigidity transition in biological tissues. Nature Physics 11, 1074–1079. doi:10.1038/nphys3471.

Bidan, C., Kommareddy, K., Rumpler, M., Kollmannsberger, P., Bréchet, Y., Fratzl, P., Dunlop, J., 2012. How linear tension converts to curvature: Geometric control of bone tissue growth. PLOS One 7, e36336. doi:10.1371/journal.pone.0036336.

Bidan, C.M., Kollmannsberger, P., Gering, V., Ehrig, S., Joly, P., Petersen, A., Vogel, V., Fratzl, P., Dunlop, J.W.C., 2016. Gradual conversion of cellular stress patterns into pre-stressed matrix architecture during in vitro tissue growth. Journal of The Royal Society Interface 13, 20160136. doi:10.1098/rsif.2016.0136.

Bidan, C.M., Kommareddy, K.P., Rumpler, M., Kollmannsberger, P., Fratzl, P., Dunlop, J.W.C., 2013. Geometry as a factor for tissue growth. Advanced Healthcare Materials 2, 186–194. doi:10.1002/adhm.201200159.

Buenzli, P., Simpson, M., 2022. Curvature dependences of wave propagation in reaction– diffusion models. Proceedings of the Royal Society A 478. doi:10.1098/rspa.2022.0582.

Buenzli, P.R., 2015. Osteocytes as a record of bone formation dynamics. Journal of Theoretical Biology 364, 418–427. doi:10.1016/j.jtbi.2014.09.028.

Buenzli, P.R., Kuba, S., Murphy, R.J., Simpson, M.J., 2025. Mechanical cell interactions on curved interfaces. Bulletin of Mathematical Biology 87. doi:10.1007/s11538-024-01406-w.

Buenzli, P.R., Lanaro, M., Wong, C.S., McLaughlin, M.P., Allenby, M.C., Woodruff, M.A., Simpson, M.J., 2020. Cell proliferation and migration explain pore bridging dynamics in 3D printed scaffolds of different pore size. Acta Biomaterialia 114, 285–295. doi:10.1016/j.actbio.2020.07.010.

Callens, S.J.P., Uyttendaele, R.J.C., Fratila-Apachitei, L.E., Zadpoor, A.A., 2020. Substrate curvature as a cue to guide spatiotemporal cell and tissue organization. Biomaterials 232, 119739. doi:10.1016/j.biomaterials.2019.119739.

Chen, C.S., Tan, J., Tien, J., 2004. Mechanotransduction at cell-matrix and cell-cell contacts. Annual Review of Biomedical Engineering 6, 275–302. doi:10.1146/annurev.bioeng.6.040803.140040.

Crandall, M.G., Lions, P.L., 1983. Viscosity solutions of Hamilton–Jacobi equations. Transactions of the American Mathematical Society 277, 1–42. doi:10.1090/S0002-9947-1983-0690039-8.

Devlin, B.L., Allenby, M.C., Ren, J., Pickering, E., Klein, T.J., Paxton, N.C., Woodruff, M.A., 2024. Materials design innovations in optimizing cellular behavior on melt electrowritten (mew) scaffolds. Advanced Functional Materials 34. doi:10.1002/adfm.202313092.

Discher, D.E., Janmey, P., Wang, Y.l., 2005. Tissue cells feel and respond to the stiffness of their substrate. Science 310, 1139–1143. doi:10.1126/science.1116995.

Dunlop, J.W.C., Fischer, F.D., Gamsjäger, E., Fratzl, P., 2010. A theoretical model for tissue growth in confined geometries. Journal of the Mechanics and Physics of Solids 58, 1073–1087. doi:10.1016/j.jmps.2010.04.008.

Dzobo, K., Thomford, N., Senthebane, D., Shipanga, H., Rowe, A., Dandara, C., Pillay, M., Motaung, K., 2018. Advances in regenerative medicine and tissue engineering: Innovation and transformation of medicine. Stem Cells International 2018, 2495848.

Ehrig, S., Schamberger, B., Bidan, C.M., West, A., Jacobi, C., Lam, K., Kollmannsberger, P., Petersen, A., Tomancak, P., Kommareddy, K., Fischer, F.D., Fratzl, P., Dunlop, J.W.C., 2019. Surface tension determines tissue shape and growth kinetics. Science Advances 5, eaav9394. doi:10.1126/sciadv.aav9394.

Eyckmans, J., Boudou, T., Yu, X., Cehn, C.S., 2011. A hitchhiker’s guide to mechanobiology. Developmental Cell 21, 35–47. doi:10.1016/j.devcel.2011.06.015.

Fozard, J.A., Byrne, H.M., Jensen, O.E., King, J.R., 2010. Continuum approximations of individual-based models for epithelial monolayers. Mathematical Medicine and Biology 27, 39–74. doi:10.1093/imammb/dqp015.

Fratzl, P., Fischer, F.D., Zickler, G.A., Dunlop, J.W.C., 2023. On shape forming by contractile filaments in the surface of growing tissues. PNAS Nexus 2. doi:10.1093/pnasnexus/pgac292.

Gardel, M.L., Shin, J.H., Mahadevan, L., Matsudaira, P., Weitz, D.A., 2004. Elastic behavior of cross-linked and bundled actin networks. Science 304, 1301–1305. doi:10.1126/science.1095087.

Ghaffarizadeh, A., Heiland, R., Friedman, S.H., Mumenthaler, S.M., Macklin, P., 2018. Physicell: An open source physics-based cell simulator for 3-d multicellular systems. PLoS Comput Biol 14, e1005991. doi:10.1371/journal.pcbi.1005991.

Goriely, A., 2017. The Mathematics and Mechanics of Biological Growth. Springer, New York, NY.

Grayson, M.A., 1987. The heat equation shrinks embedded plane curves to round points. Journal of Differential Geometry 26, 285–314. doi:10.4310/jdg/1214441371.

Guyot, Y., Papantoniou, I., Chai, Y.C., Van Bael, S., Schrooten, J., Geris, L., 2014. A computational model for cell/ecm growth on 3D surfaces using the level set method: a bone tissue engineering case study. Biomechanics and Modeling in Mechanobiology 13, 1361–1371. doi:10.1007/s10237-014-0577-5.

Han, W., He, W., Yang, W., Li, J., Yang, Z., Lu, X., Qin, A., Qian, Y., 2016. The osteogenic potential of human bone callus. Scientific Reports 6. doi:10.1038/srep36330.

Hegarty-Cremer, S.G.D., Simpson, M.J., Andersen, T.L., Buenzli, P.R., 2021. Modelling cell guidance and curvature control in evolving biological tissues. Journal of Theoretical Biology 520, 110658. doi:10.1016/j.jtbi.2021.110658.

Hollister, S.J., 2005. Porous scaffold design for tissue engineering. Natural Materials 4, 518–524. doi:10.1038/nmat1421.

Jansen, K.A., Donato, D.M., Balcioglu, H.E., Schmidt, T., Danen, E.H., Koenderink, G.H., 2015. A guide to mechanobiology: Where biology and physics meet. Biochimica et Biophysica Acta (BBA) - Molecular Cell Research 1853, 3043–3052. doi:10.1016/j.bbamcr.2015.05.007.

Jones, G., Chapman, S., 2012. Modeling growth in biological materials. SIAM Review 54, 52–118. doi:10.1137/080731785.

Kim, S., Pochitaloff, M., Stooke-Vaughan, G.A., Campàs, O., 2021. Embryonic tissues as active foams. Nature Physics 17, 859–866. doi:10.1038/s41567-021-01215-1.

Kreutz, C., Raue, A., Kaschek, D., Timmer, J., 2013. Profile likelihood in systems biology. The FEBS Journal 280, 2564–2571. doi:10.1111/febs.12276.

Kuba, S., Buenzli, P.R., Simpson, M.J., 2026. Curvature controlled tissue growth incorporating mechanical cell interactions. URL: https://github.com/Shahak-Kuba/Kuba2024_MechanicalCell_CurvatureControl_Growth.

Kurganov, A., Tadmor, E., 2000. New high-resolution central schemes for nonlinear conservation laws and convection–diffusion equations. Journal of Computational Physics 160, 241–282. doi:10.1006/jcph.2000.6459.

Ladoux, B., Mège, R.M., 2017. Mechanobiology of collective cell behaviours. Nature Reviews Molecular Cell Biology 18, 743–757. doi:10.1038/nrm.2017.98.

Lanaro, M., Mclaughlin, M.P., Simpson, M.J., Buenzli, P.R., Wong, C.S., Allenby, M.C., Woodruff, M.A., 2021. A quantitative analysis of cell bridging kinetics on a scaffold using computer vision algorithms. Acta Biomaterialia 136, 429–440. doi:10.1016/j.actbio.2021.09.042.

Lee, H., Park, J., Yoon, S., Lee, C., Kim, J., 2019. Mathematical model and numerical simulation for tissue growth on bioscaffolds. Applied Sciences 9, 4058. doi:10.3390/app9194058.

Lee, Y., Lim, W., 2017. Shoelace formula: Connecting the area of a polygon and the vector cross product. The Mathematics Teacher 110, 631–636. doi:10.5951/mathteacher.110.8.0631.

LeVeque, R.J., 2002. Finite Volume Methods for Hyperbolic Problems. Cambridge University Press. doi:10.1017/CBO9780511791253.

Lim, C.T., Bershadsky, A., Sheetz, M.P., 2010. Mechanobiology. Journal of The Royal Society Interface 7, S291–S293. doi:10.1098/rsif.2010.0150.

Marotti, G., Zallone, A., Ledda, M., 1975. Number, size and arrangement of osteoblasts in osteons at different stages of formation, in: Calcified Tissues 1975, pp. 96–101. doi:10.1007/978-3-662-29272-3_13.

Moeendarbary, E., Harris, A.R., 2014. Cell mechanics: principles, practices, and prospects. WIREs Systems Biology and Medicine 6, 371–388. doi:10.1002/wsbm.1275.

Murphy, R.J., Buenzli, P.R., Baker, R.E., Simpson, M.J., 2019. A one-dimensional individual-based mechanical model of cell movement in heterogeneous tissues and its coarse-grained approximation. Proceedings of the Royal Society A 475, 20180838. doi:10.1098/rspa.2018.0838.

Murphy, R.J., Maclaren, O.J., Simpson, M.J., 2024. Implementing measurement error models with mechanistic mathematical models in a likelihood-based framework for estimation, identifiability analysis and prediction in the life sciences. Journal of The Royal Society Interface 21. doi:10.1098/rsif.2023.0402.

Murray, P.J., Edwards, C.M., Tindall, M.J., Maini, P.K., 2009. From a discrete to a continuum model of cell dynamics in one dimension. Physical Review E 80, 031912. doi:10.1103/PhysRevE.80.031912.

Nelson, C.M., 2022. Mechanical control of cell differentiation. Annual Review of Biomedical Engineering 24, 307–322. doi:10.1146/annurev-bioeng-060418-052527.

Nelson, C.M., Jean, R.P., Tan, J.L., Liu, W.F., Sniadecki, N.J., Spector, A.A., Chen, C.S., 2005. Emergent patterns of growth controlled by multicellular form and mechanics. Proceedings of the National Academy of Sciences 102, 11594–11599. doi:10.1073/pnas.0502575102.

O’Brien, F.J., 2011. Biomaterials & scaffolds for tissue engineering. Materials Today 14, 88–95. doi:10.1016/S1369-7021(11)70058-X.

Osborne, J.M., Fletcher, A.G., Pitt-Francis, J.M., Maini, P.K., Gavaghan, D.J., 2017. Comparing individual-based approaches to modelling the self-organization of multicellular tissues. PLoS Comput Biol 13, e1005387. doi:10.1371/journal.pcbi.1005387.

Qiu, Z.Y., Cui, Y., Wang, X.M., 2019. Mineralized collagen bone graft substitutes, Woodhead Publishing, pp. 1–22. doi:10.1016/B978-0-08-102717-2.00001-1.

Rackauckas, C., Nie, Q., 2017. DifferentialEquations.jl — A performant and feature-rich ecosystem for solving differential equations in julia. URL: https://github.com/JuliaDiffEq/DifferentialEquations.jl.

Riccobelli, D., 2025. Surface tension-driven boundary growth in tumour spheroids. Interface Focus 15. doi:10.1098/rsfs.2024.0035.

Ripamonti, U., Roden, L., 2010. Biomimetics for the induction of bone formation. Expert Review of Medical Devices 7, 469–479. doi:10.1586/erd.10.17.

Rodriguez, E., Hoger, A., McCulloch, A., 1994. Stress-dependent finite growth in soft elastic tissues. Journal of Biomechanics 27, 455–467. doi:10.1016/0021-9290(94)90021-3.

Rumpler, M., Woesz, A., Dunlop, J.W., van Dongen, J.T., Fratzl, P., 2008. The effect of geometry on three-dimensional tissue growth. Journal of The Royal Society Interface 5, 1173–1180. doi:10.1098/rsif.2008.0064.

Schamberger, B., Ziege, R., Anselme, K., Ben Amar, M., Bykowski, M., Castro, A.P.G., Cipitria, A., Coles, R.A., Dimova, R., Eder, M., Ehrig, S., Escudero, L.M., Evans, M.E., Fernandes, P.R., Fratzl, P., Geris, L., Gierlinger, N., Hannezo, E., Iglič, A., Kirkensgaard, J.J.K., Kollmannsberger, P., Kowalewska, Ł., Kurniawan, N.A., Papantoniou, I., Pieuchot, L., Pires, T.H.V., Renner, L.D., Sageman-Furnas, A.O., Schröder-Turk, G.E., Sengupta, A., aSharma, V.R., Tagua, A., Tomba, C., Trepat, X., Waters, S.L., Yeo, E.F., Roschger, A., Bidan, C.M., Dunlop, J.W.C., 2023. Curvature in biological systems: Its quantification, emergence, and implications across the scales. Advanced Materials 35. doi:10.1002/adma.202206110.

Sethian, J.A., 1999. Level Set Methods and Fast Marching Methods: Evolving Interfaces in Computational Geometry, Fluid Mechanics, Computer Vision, and Materials Science. Cambridge University Press.

Simpson, M.J., Baker, R.E., 2026. Parameter identifiability, parameter estimation and model prediction for differential equation models. SIAM Review 68, 153–171. doi:10.1137/24M1667968.

Simpson, M.J., Landman, K.A., Hughes, B.D., 2010. Cell invasion with proliferation mechanisms motivated by time-lapse data. Physica A: Statistical Mechanics and its Applications 389, 3779–3790. doi:10.1016/j.physa.2010.05.020.

Simpson, M.J., Treloar, K.K., Binder, B.J., Haridas, P., Manton, K.J., Leavesley, D.I., McElwain, D.L.S., Baker, R.E., 2013. Quantifying the roles of cell motility and cell proliferation in a circular barrier assay. Journal of The Royal Society Interface 10, 20130007. doi:10.1098/rsif.2013.0007.

Storm, C., Pastore, J.J., MacKintosh, F.C., Lubensky, T.C., Janmey, P.A., 2005. Nonlinear elasticity in biological gels. Nature 435, 191–194. doi:10.1038/nature03521.

Taber, L., 1995. Biomechanics of growth, remodeling and morphogenesis. Applied Mechanics Reviews 48, 487–545.

Taber, L.A., 2020. Continuum modeling in mechanobiology. Springer International Publishing, Switzerland.

Tambyah, T.A., Murphy, R.J., Buenzli, P.R., Simpson, M.J., 2020. A free boundary mechanobiological model of epithelial tissues. Proceedings of the Royal Society A 476. doi:10.1098/rspa.2020.0528.

VandenHeuvel, D.J., Devlin, B.L., Buenzli, P.R., Woodruff, M.A., Simpson, M.J., 2023. New computational tools and experiments reveal how geometry affects tissue growth in 3D printed scaffolds. Chemical Engineering Journal 475, 145776. doi:10.1016/j.cej.2023.145776.

Vetter, R., Runser, S.V.M., Iber, D., 2024. PolyHoop: Soft particle and tissue dynamics with topological transitions. Computer Physics Communications 299. doi:10.1016/j.cpc.2024.109128.

Wang, J., Thampatty, B., 2006. An introductory review of cell mechanobiology. Biomechanics and Modeling in Mechanobiology 5, 1–16. doi:10.1007/s10237-005-0012-z.

Wozniak, P., El Haj, A.J., 2007. 14 - bone regeneration and repair using tissue engineering, in: Boccaccini, A.R., Gough, J.E. (Eds.), Tissue Engineering Using Ceramics and Polymers. Woodhead Publishing, Philadelphia. Woodhead Publishing Series in Biomaterials, pp. 294–318. doi:10.1533/9781845693817.2.294.

Xi, W., Saw, T.B., Delacour, D., Lim, C.T., Ladoux, B., 2019. Material approaches to active tissue mechanics. Nature Reviews Materials 4, 23–44. doi:10.1038/s41578-018-0066-z.

